# Genome-Scale Characterization of Predicted Plastid-Targeted Proteomes in Higher Plants

**DOI:** 10.1101/867242

**Authors:** Ryan W. Christian, Seanna L. Hewitt, Eric H. Roalson, Amit Dhingra

**Author notes:** Corresponding author Email Addresses: RWC, SLH, EHR, AD.

## Abstract

Plastids are morphologically and functionally diverse organelles that are dependent on nuclear-encoded, plastid-targeted proteins for all biochemical and regulatory functions. However, how plastid proteomes vary temporally, spatially, and taxonomically has been historically difficult to analyze at genome-wide scale using experimental methods. A bioinformatics workflow was developed and evaluated using a combination of fast and user-friendly subcellular prediction programs to maximize performance and accuracy for chloroplast transit peptides and demonstrate this technique on the predicted proteomes of 15 sequenced plant genomes. Gene family grouping was then performed in parallel using modified approaches of reciprocal best BLAST hits (RBH) and UCLUST. Between 628 protein families were found to have conserved plastid targeting across angiosperm species using RBH, and 828 using UCLUST. However, thousands of clusters were also detected where only one species had predicted plastid targeting, most notably in *Panicum virgatum* which had 1,458 proteins with species-unique targeting. An average of 45% overlap was found in plastid-targeted gene families compared with Arabidopsis, but an additional 20% of proteins matched against the full Arabidopsis proteome, indicating a unique evolution of plastid targeting. Neofunctionalization through subcellular relocalization is known to impart novel biological functions but has not been described before on genome-wide scale for the plastid proteome. Further work to correlate these predicted novel plastid-targeted proteins to transcript abundance and high-throughput proteomics will uncover unique aspects of plastid biology and shed light on how the plastid proteome has evolved to change plastid morphology and biochemistry.

## Introduction

Plastids represent biochemically and morphologically complex organelles and can change both form and function drastically in response to developmental and environmental cues. A vestigial but functional genome of 120-160 kb harboring ∼90 protein-coding genes is present in the plastids of photosynthetic higher plants [1]. However, the total chloroplast proteome conservatively contains 2,000-3,500 proteins as reported in Arabidopsis [2–4], but as many as 4,875 plastid-targeted proteins are estimated in eSLDB [5], and 5,136 by the Chloroplast 2010 project [6–8]. Less than 900 of 4,500 genes horizontally transferred from the ancestral cyanobacterium are predicted to be retargeted to the plastid *in vivo* [9].

There seems to be a difference between the composition of plastid-targeted proteomes in dicots and monocots. Only 21% of plastid-targeted rice proteins have a predicted homolog in the predicted Arabidopsis plastid proteome, and in reciprocal comparison the number is 38% [2]. A similar result was obtained in a comparison of six crop plants against Arabidopsis, in which an average of 51.0% of the predicted plastid proteome of each species matched sequences to the Arabidopsis predicted plastid proteome, while 67.5% matched against the full Arabidopsis proteome [10]. Thus, the plastid pan-proteome is extremely diverse and is composed of unique proteins at the species-level. Furthermore, as the number of conserved sequences across all the genomes analyzed closely mirrors the number of genes of cyanobacterial origin, the non-conserved plastid-targeted genes most likely evolved from eukaryotic sequences. The variability in the predicted plastid proteome mirrors the observable diversity in plastid function and ultrastructure in different species or under different environmental and developmental conditions [2,10–13]. The diversity of plastid proteomes is evident even within the same plastid morphotype: the pigment-storing chromoplast alone has at least four described ultrastructural phenotypes across various species with unique sub-organellar membrane structures that can occur either singly or mixed within individual plastids [14]. Morphological differences in plastid shape and ultrastructure are noted even in genetically similar cultivars of the same species. Both chloroplasts and chromoplasts of developing apple peel differ significantly from tomato, which is used as a model reference for chromoplast differentiation in fruits [15, 16]. Variation has also been documented between the apple cultivars and the epidermal and collenchymal plastids [11].

The observed phenotypic diversity of plastids could be explained by three potential molecular factors: 1.) Differences in the expression of genes controlling the rate and total amount of protein accumulation or import. This aspect could lead to unique phenotypes without necessarily changing the subset of plastid-targeted proteins. 2.) Mutations within a shared group of plastid-targeted proteins could lead to neofunctionalization. 3.) Finally, gain or loss of transit peptides causing subcellular mistargeting could alter the total pool of plastid-targeted proteins.

These factors are not mutually exclusive, and examples of each mechanism are known. Gene expression differences, possibly caused by epigenetic DNA methylation patterns, are responsible for differential protein accumulation in mesophyll and bundle sheath cells of C4 plants, illustrating the first point [17–20]. In support of the second mechanism, point mutations in the active site of plastid-targeted limonene synthase change the abundance and distribution of different monoterpenoid end products in bacterial expression systems [21], and transplastomic expression of a delta-9 desaturase gene causes changes in fatty acid concentrations and levels of unsaturation, cold tolerance, leaf senescence, and seed yield [22] are additional examples. While it is challenging to address the neofunctionalization of plastid-targeted proteins via mutation without detailed reverse genetics experiments, the other mechanisms can be evaluated with high-throughput sequencing and bioinformatics.

High-throughput proteomics using mass spectrometry (MS) has been an important means of surveying organellar proteomes and comprises the majority of current plastid proteome evidence. However, these techniques have historically been limited to the chloroplast morphotype and a restricted number of plant species. Excellent databases for high-throughput plastid proteomes based largely on mass spectrometry are accessible at AT_CHLORO [23], PPDB [24], SUBA4 [25], and CROPPAL [26]. However, caution should be exercised in interpreting these datasets because MS is susceptible to high false positive errors due to contamination during plastid isolation, liberal mass tolerance, and errors in peptide mapping, among other problems [27–29]. While the use of reference genomes and transcriptomes can help overcome peptide mapping issues, other technical issues are more difficult to resolve. Use of fluorescent protein chimeras (e.g., GFP – green fluorescent protein), though lower-throughput, typically have higher biological accuracy. Using these, localization of low-abundance, as well as proteins from species lacking robust plastid isolation methods, can be evaluated with higher efficiency. However, GFP techniques are not immune to experimental error either. Since the sequence of the mature protein partially influences localization (e.g., [30–32]), GFP fused to the native protein may alter localization in some cases. Furthermore, dual-targeted mitochondrial/chloroplast proteins can be mislocalized in GFP assays [33]. Alternative transcripts or alternative protein products may also produce differential subcellular localization that are either not captured in GFP assays or give ambiguous results. Given these experimental limitations, a robust bioinformatics workflow could enable rapid and cost-effective assessment of plastid proteomes with somewhat comparable accuracy. Though wet lab validation is still necessary, these datasets could narrow the focus to smaller subsets of proteins of interest which could be more manageably targeted for wet lab validation depending on the biological question being asked.

The semi-conserved and sometimes ambiguous nature of chloroplast transit peptides makes *in silico* predictions challenging. However, sequence- and annotation-based approaches have yielded results with significant accuracy. Protein sequence-based prediction uses the amino acid content or the presence of conserved motifs in the peptide to make predictions. Use of the amino acid content alone, such as in the tool PCLR, is enough to predict many plastid-targeted proteins [34]. More complex sequence-based identify conserved motifs, such as in iPSORT [35] and WoLF-PSORT [36], or sliding-window searching algorithm such as Localizer [37], make predictions based on the sum of prediction vectors to determine transit peptide similarity. Finally, tools that use neural networks such as ChloroP [38], TargetP [39, 40], Predotar [41], PredSL [42], and Protein Prowler [43] use multiple layers of nodes to identify the best-scoring localization. In contrast, annotation-based methods such as CLPFD [44] and EpiLoc [45], or simple text-based methods based on GO annotations [46], use homology to proteins with known localization to designate subcellular predictions. While these methods offer advantages over sequence-based methods for proteins with annotated homologs, they perform poorly for novel proteins [47]. Hybrid approaches including MultiLoc2 [48], Sherloc2 [49], Y-Loc [50], and Plant-mPLoc [51] combine sequence- and annotation-based methods in an attempt to overcome this limitation. Unfortunately, the homology component of hybrid approaches is weighted more heavily, which can lead to the false prediction of proteins with transit peptide variation or for proteins with shared domains. Both high-throughput proteomics and bioinformatics approaches consistently indicate that the plastid proteome content is highly dynamic and likely has significant variability across the plant kingdom. With newer methods, ever-growing genomic resources, and availability of better gene annotation methods, earlier estimates of conserved and non-conserved sets of the plastid proteome warrant an update.

This study evaluated the hypothesis that bioinformatics methods could achieve similar accuracy to experimental methods by comprehensively testing previously published subcellular prediction algorithms both alone and in combination. A specific combination of methods was found to be most efficient, which was then used to globally predict nuclear-encoded plastid-targeted proteins for fifteen higher plant species including eight eudicots, six monocots, and *Amborella trichopoda*, an early diverging species of the angiosperm clade. Two parallel approaches, Reciprocal-Best Blast Hit (RBH) and UCLUST (Edgar, 2010) were used to perform clustering, and the sub-cellular localization prediction for each cluster was analyzed to identify conserved, semi-conserved, and non-conserved plastid-targeted proteins. This approach evaluated the hypothesis that a relative minority of plastid-targeted genes are conserved among all species. It was found that natural selection and environmental influence has shaped the development of species-specific plastid proteomes.

## Results and Discussion

### Identification of Optimal Subcellular Prediction Workflows

To test the hypothesis that a bioinformatics workflow could reach parity with experimental methodology, the accuracy of six subcellular prediction algorithms including TargetP [39], WoLF PSORT [36], PredSL [42], Localizer [37], Multiloc2 [48], and PCLR [34] was first evaluated using data from the original publications. Sensitivity, specificity, accuracy, and Matthew’s Correlation Coefficient (MCC) were evaluated for each program as it related to the prediction of plastid-targeted proteins (Table 1). Sensitivity, specificity, and MCC in TargetP were found to exactly match the values reported by Emanuelsson et al. (2000, 2007) and while minor differences were found for MultiLoc and PredSL, these discrepancies likely represent rounding errors. Unexpectedly, significant differences were found for PCLR and Localizer: in PCLR, sensitivity was found to be 52.1%, which was about 5% lower than what was reported [34]. In Localizer, calculated specificity was 78.9%, nearly 16% lower than the 95.7% reported [37]. In both cases, all other performance statistics were identical or nearly identical, so it is likely that the discrepancies in Localizer and PCLR represent either miscalculations or transcriptional errors in the data used for analysis in the original publications.

**Table 1:** Self-Reported Performance of Six Algorithms on Prediction of Plastid-Targeted Proteins. . Self-reported values for overall and plastidial sensitivity (SE), specificity (SP), Matthew’s Correlation Coefficient (MCC), and accuracy (ACC). Parentheses indicate values that were calculated to be different from the original paper using the same data. Programs marked with an asterisk (*) had a confusion matrix available, while those marked with a cross (†) did not, but confusion matrices were inferred by the available data; estimations were left as non-integer values, and therefore suffer from rounding errors in MCC and ACC calculations. Localizer, marked with a double cross (‡), was re-run with the original dataset provided in the publication’s supplementary information.

| Algorithm | Source | Training Dataset(s) | # of Training Sequences | Plastidial SE | Plastidial SP | Plastidial MCC | Plastidial ACC |
| --- | --- | --- | --- | --- | --- | --- | --- |
| **TargetP\*** | [39,40] | SWISS-PROT releases 36,37,38 | 940 | 0.85 | 0.69 | 0.72 | N/A (0.921) |
| **WolfPSORT** | [36] | Uniprot version 45 | 2,113 | 0.7 | 0.7 | N/A | N/A |
| **PredSL†** | [42] | Various (Uniprot release 3.5) | 1,002 | 0.9 | 0.91 | 0.88 (0.874) | N/A |
| **Localizer‡** | [37] | CropPAL (GFP only) | 410 | 0.725 | 0.957 (0.798) | 0.71 | 0.914 (0.916) |
| **Multiloc2 (Low-Res)†** | [48] | BaCelLo Independent Dataset | 132 | 0.77 | 0.53 | 0.72 | N/A (0.853) |
| **Multiloc2 (High-Res)†** | [48] | BaCelLo Independent Dataset | 132 | 0.53 | 0.94 | 0.51 (0.539) | N/A (0.735) |
| **PCLR\*** | [34] | ChloroP, TargetP | 847 | 0.87 (0.821) | 0.30 (0.301) | 0.372 | 0.720 |

Next, cross-validation of subcellular prediction programs was performed against proteins with experimentally-determined subcellular localization retrieved from AT_CHLORO [23], PPDB [24, 52], CropPAL and CropPAL2 [26] and Suba4 [25,53–55], resulting in 42,761 nonredundant sequences including 32,450 proteins validated by mass spectrometry (MS) and 3,722 validated by GFP. Most prediction algorithms were found to have lower performance against biological data than reported in the original reports, as shown in Table 2 and Figure 1.

**Figure 1:**
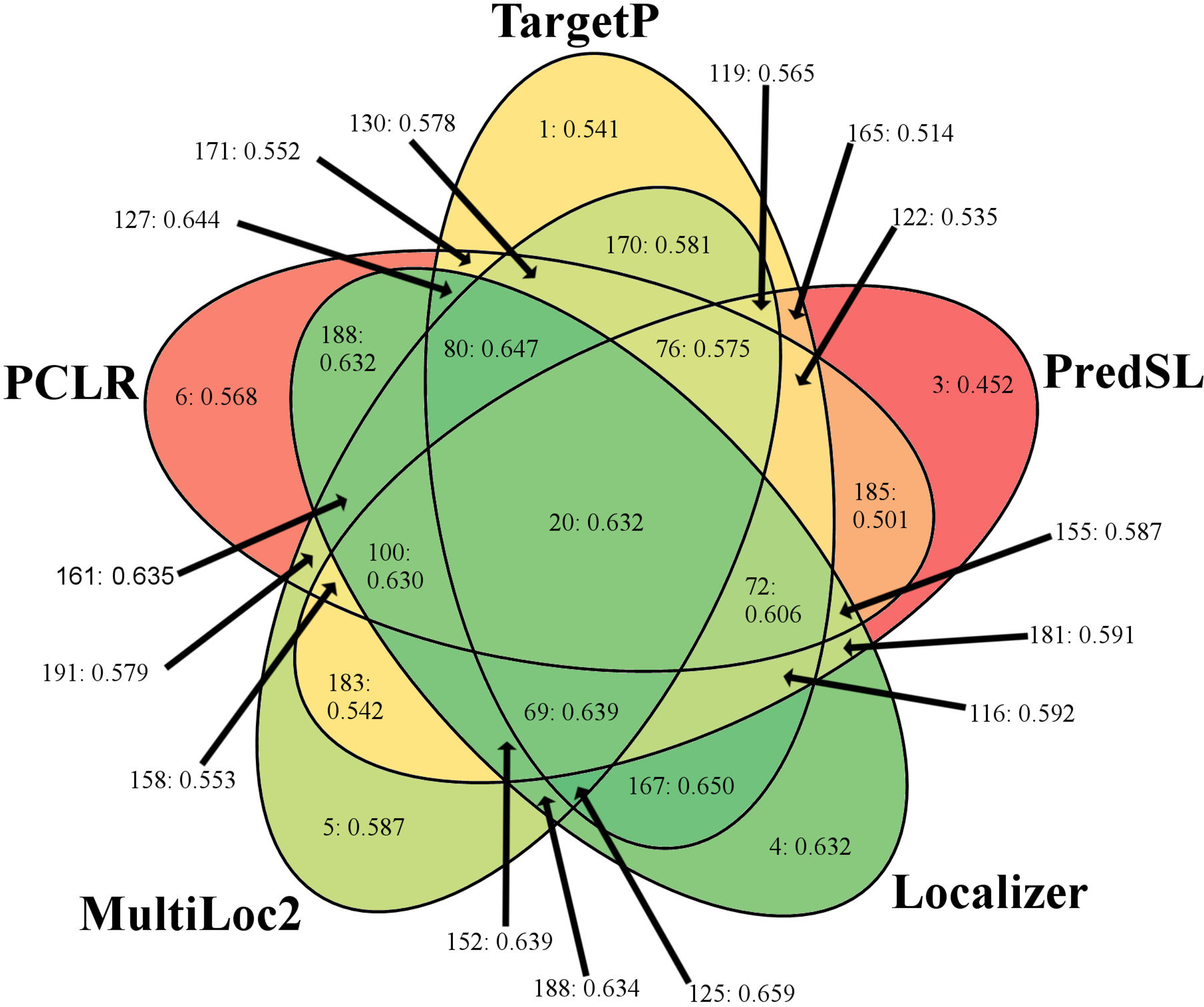
Venn-Diagram of Combinatorial and Standalone Subcellular Prediction Algorithms. Performance measured by MCC on proteins with subcellular localization validated by GFP is represented as a heatmap with high values in green and low values in red. For each intersection, only the best accept threshold is represented. Numbers indicate workflow number followed by the calculated MCC.

**Table 2:**
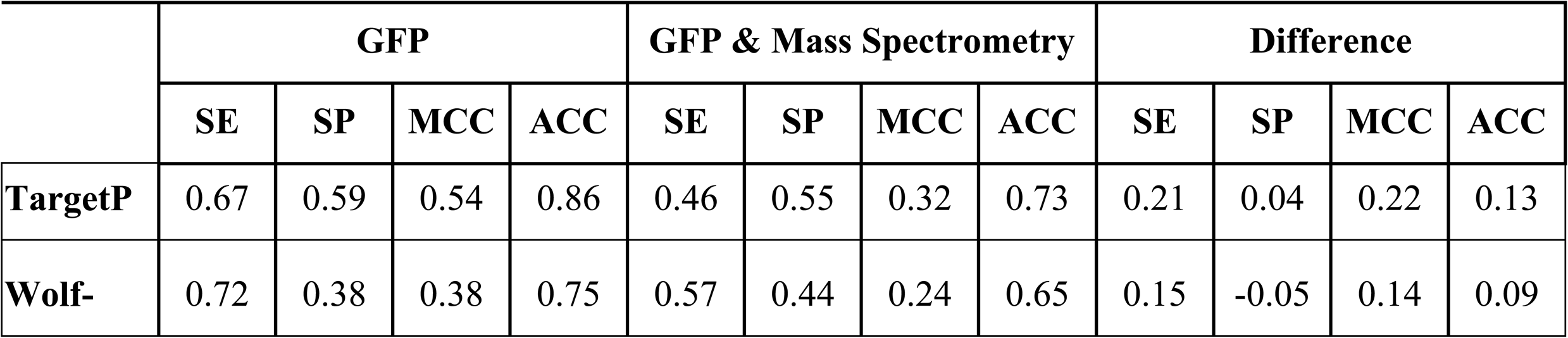
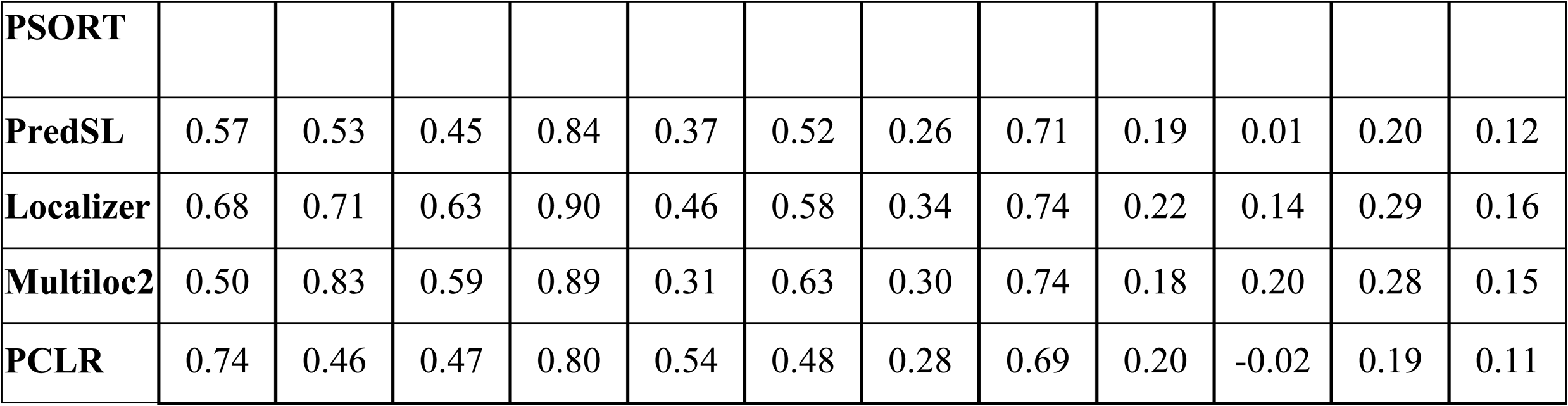
Review of Algorithms using modern curated datasets (combined) . For each program, SE, SP, MCC, and ACC are reported compared to *in vivo* experimental data using a conservative dataset of GFP-validated proteins, or a larger but more liberal dataset comprised of both GFP and MS data. Difference between observed performance statistics of different datasets is presented as GFP minus MS/GFP. MS data was found to have increased error especially for observed sensitivity, indicating that a large number of MS-validated proteins are likely artefactual. Furthermore, this suggests that the overall performance of subcellular prediction methods is likely more accurate than high-throughput proteomics papers suggest.

However, substantial differences were observed based on the method of experimental validation. On average among the six algorithms, sensitivity was 15.7% higher in the GFP-validated dataset while no significant change in specificity was found; this difference resulted in 10% higher overall accuracy and an increase of 0.159 in MCC for GFP-validated proteins. By further narrowing focus to a dataset of proteins validated by both methods, sensitivity increased by an additional 7.6%, and specificity increased 2.5%, on average. Due to the previously reported high false positive rates associated with shotgun proteomics of organellar proteomes [27, 28], program performance was expected to be much higher for GFP-validated proteins. While the dataset containing proteins experimentally validated by both GFP and mass spectrometry showed the highest apparent performance for the six subcellular prediction algorithms - and is likely closer to the biological accuracy of these programs - it contains roughly a third as many proteins as the GFP-validated dataset and is heavily biased by Arabidopsis sequences. Therefore, remaining comparisons focused on the GFP-validated dataset. Similarly, MCC was used as the primary measure of biological accuracy of *in silico* approaches to avoid problems due to drastically different dataset sizes.

Overall, the highest-performing program in terms of MCC was Localizer, followed by MultiLoc2-HR, TargetP, PCLR, PredSL, WoLF PSORT, and MultiLoc2-LR. Of these, PredSL and MultiLoc2-LR performed poorly with GFP-validated proteins compared to the original reports, while other programs decreased marginally or performed similarly to the published MCC. Among the six programs that were evaluated, Localizer had the highest performance regardless of the experimental method used for validation, which is surprising since it is a simpler tool than annotation-based methods which have been at the forefront of subcellular prediction methods recently. Part of Localizer’s increased accuracy may be due to its unique capacity to predict dual-targeted mitochondrial/chloroplast proteins. Over 200 dual-localized proteins have been described in Arabidopsis [56] and over 500 are predicted to have ambiguous transit peptides [57]. Increased accuracy in the prediction of these sequences in Localizer could alone account for a portion of its higher performance. After Localizer, MultiLoc2 had the next-highest MCC and also had the highest specificity of any program, at 83% in GFP-validated proteins. MultiLoc is a hybrid method combining annotation and sequence analysis, so these findings support that the use of hybrid methods yields robust biological specificity. However, MultiLoc also had the worst sensitivity of any program, correctly predicting only 50% of bonafide plastid-targeted proteins validated by GFP or 31% of sequences validated by either GFP or mass spectrometry. TargetP, which has historically been the most popular subcellular prediction program for plants since its introduction, was found to perform at lower accuracy than earlier estimates: even when using the more conservative GFP-validated data, specificity was only 59% and sensitivity was 67%. Previous experiments using high-throughput shotgun proteomics have reported that the sensitivity of TargetP is as low as 62% [3,58–60]. Use of strictly-curated data improves the apparent sensitivity up to 86%, but false positive rates are still problematic as a specificity of about 65% is observed [61]. The results presented here suggest that the biological accuracy of TargetP is somewhat closer to the initial estimates on non-curated data. PredSL, PCLR, and WoLF-PSORT were the lowest-ranked programs by MCC for prediction of plastid-targeted proteins, in that order, but typically had higher sensitivity than Localizer or MultiLoc2.

Differences in the amino acid composition of transit peptides are observable between rice and Arabidopsis, which have an overrepresentation of alanine and serine, respectively [61]. Therefore, differences in the prediction of monocot or eudicot sequences were assessed, and different programs displayed significant bias (Table 3). PCLR was the most drastically affected, with an MCC bias of +0.091 in monocots, representing a roughly 20% increase compared with eudicots. This finding is somewhat unsurprising because PCLR is the only program which uses sequence composition alone to make predictions and is, therefore, more susceptible to bias than motif-or annotation-based methods. TargetP was the only other tool that favored monocots, with an increase of 0.055 (+10.2%) in MCC. A marginal difference between monocot and eudicot prediction was observed when Localizer was used, which differed by only 0.008 in MCC, slightly favoring eudicots. Eudicot sequences were favored in the other prediction programs, with between 0.043 (+10%) higher MCC in WoLF POSRT and 0.066 in PredSL (+14.9%). To the best of our knowledge, this is the first study to report this type of error or bias for *in silico* prediction methods. Some differences have also been described for the proposed subunits of the TIC translocon in grasses, which could result in coevolution of the transit peptide sequence composition [62–64]. Choice of training and cross-validated datasets could significantly sway the predictions of sequence-based methods, while overrepresentation or prioritization of sequences for Arabidopsis and thereby eudicots could introduce bias to annotation-based methods. Although these species-specific differences are smaller than differences observed for sequences validated by mass spectrometry compared with GFP, they are still noteworthy and have consequences for whole-genome prediction. In contrast, WoLF-PSORT and Localizer were found to have insignificant if any bias, making them attractive both as standalone programs or in combinatorial approaches where they could mask biases of other programs.

**Table 3:** Performance of prediction algorithms against GFP-validated proteins from monocots and eudicots. . Performance of each prediction algorithm in monocots and eudicots and the difference between these datasets is presented; dataset sizes are roughly similar for monocot and eudicot sequences, but MCC is still preferable for comparison. 161 plastid-localized proteins and 640 non-plastid-targeted proteins are included for monocots, while eudicots include 489 plastid-targeted and 2,432 non-plastid-targeted proteins.

|  | Monocot: GFP |  |  |  | Eudicot: GFP |  |  |  | Monocot-Eudicot |  |  |  |
| --- | --- | --- | --- | --- | --- | --- | --- | --- | --- | --- | --- | --- |
|  | SE | SP | MCC | ACC | SE | SP | MCC | ACC | SE | SP | MCC | ACC |
| **TargetP** | 0.62 | 0.71 | 0.59 | 0.87 | 0.68 | 0.56 | 0.53 | 0.86 | -0.06 | 0.15 | 0.05 | 0.02 |
| **Wolf-PSORT** | 0.72 | 0.38 | 0.35 | 0.71 | 0.72 | 0.38 | 0.39 | 0.76 | 0.00 | -0.01 | -0.04 | -0.05 |
| **PredSL** | 0.43 | 0.59 | 0.40 | 0.83 | 0.61 | 0.52 | 0.47 | 0.84 | -0.17 | 0.06 | -0.07 | -0.02 |
| **Localizer** | 0.63 | 0.76 | 0.63 | 0.89 | 0.69 | 0.70 | 0.63 | 0.90 | -0.06 | 0.06 | -0.01 | -0.01 |
| **Multiloc2** | 0.40 | 0.89 | 0.54 | 0.87 | 0.53 | 0.81 | 0.60 | 0.90 | -0.12 | 0.08 | -0.06 | -0.03 |
| **PCLR** | 0.75 | 0.56 | 0.54 | 0.83 | 0.73 | 0.43 | 0.45 | 0.80 | 0.01 | 0.12 | 0.09 | 0.03 |

### Combinatorial Workflow Outperforms Single Programs

Use of multiple prediction algorithms in combination is a powerful strategy to combine the strengths and overcome the limitations of single programs. Combinatorial approaches have been used to improve the accuracy of predictions in whole-genome analyses (e.g., [2]) or to curate mass spectrometry data (e.g., [65–68]). Additionally, a combinatorial workflow using 22 prediction algorithms and four experimental techniques is used in the SUBAcon algorithm implemented for the SUBA4 database of Arabidopsis proteins which reportedly yields up to 97.5% accuracy for chloroplast localization and 90% for other compartments [25, 55]. While SUBAcon does not strictly require experimental data to perform predictions, available evidence weighs heavily on the final prediction and contributes to the reported accuracy. Even if experimental evidence were to be ignored, the use of 22 separate subcellular prediction algorithms is not feasible for individual researchers or application to enormous datasets. Therefore, a bioinformatics-based workflow that can work efficiently would be desirable.

Calculations were performed for each possible permutation of subcellular prediction algorithms and for all possible acceptable thresholds for each combination as applied to GFP-validated proteins. For example, for the combination of TargetP, PredSL, and Localizer, three thresholds were tested in which one, two, or all three programs needed to predict plastid localization to consider that protein as having a plastid transit peptide. To simplify analyses, the poorly-performing WoLF PSORT was removed from consideration (results including WoLF PSORT and datasets including MS-validated proteins are available in Additional File 2). In total, 80 unique workflows including the five remaining standalone program workflows were evaluated against GFP-validated proteins, the results of which are graphically summarized in Figure 1, and numerically ranked by MCC in Table 4. Unequivocally, the results demonstrate that combinations of programs tend to outperform single programs for GFP-validated data: among the 25 workflows with the highest MCC, 23 were combinatorial approaches, while the standalone Localizer ranked tenth and Multiloc2-HR 22^nd^. Localizer was not only the best-performing standalone program but was also overrepresented in combinatorial workflows: except the standalone Multiloc2-HR workflow, Localizer appeared in all 25 top-performing workflows. It is interesting to note that combinations that rank higher tend to combine programs with high sensitivity with counterparts that have lower sensitivity but higher specificity, thus correcting for each other’s deficiencies. Specifically, most of the combinations with the highest MCC and ACC tend to include Localizer most often, followed by MultiLoc2, TargetP, PCLR, and lastly PredSL. The ranking of Localizer is unsurprising given that its relatively balanced and high sensitivity and specificity are unparalleled by any of the other programs. However, MultiLoc2’s extremely high specificity makes it a valuable component of many workflows despite its low sensitivity. The best performing workflow used TargetP, Localizer, and Multiloc2 and required 2 of the three programs to predict plastid targeting to define a sequence as containing a plastid transit peptide; specificity of 78.5%, the sensitivity of 64.6%, and MCC of 0.659 was achieved with this approach. In comparison to TargetP alone, a nearly 20% increase in specificity was observed with no loss in sensitivity. However, as the annotation-based functions of MultiLoc2 make it difficult to run on extensive datasets, an alternative workflow using a “2 of 2” consensus approach for TargetP and Localizer was found which ranked 2^nd^ and achieved a marginally higher specificity of 80.7%. Furthermore, comparing the accuracy of the best workflows to Table 2 and to prior evaluations of experimental methodology (e.g., [61]) supported the hypothesis that bioinformatics methods could reach parity with mass spectrometry in characterizing the plastid proteome. Due to the increased simplicity and comparable performance of the TargetP/Localizer consensus approach, this workflow was selected for subsequent genome-scale prediction of plastid-targeted proteins.

**Table 4:** Best combinatorial prediction approaches ranked by Matthew’s Correlation Coefficient (MCC) . The sensitivity (SE), specificity (SP), Matthew’s Correlation Coefficient (MCC), and accuracy (ACC) are presented for each program. Almost all of the highest-performing programs utilized Localizer in their approach, followed by Multiloc2 and TargetP. Localizer and MultiLoc2 were also the only two programs which ranked highly as standalone algorithms, whereas the remaining workflows used two or more individual programs.

| Rank | Workflow | Description | SE | SP | MCC | ACC |
| --- | --- | --- | --- | --- | --- | --- |
| 1 | 125 | 2/3 of (TargetP, Localizer, Multiloc2) | 0.646 | 0.785 | 0.659 | 0.907 |
| 2 | 167 | 2/2 of (TargetP, Localizer) | 0.611 | 0.807 | 0.650 | 0.907 |
| 3 | 80 | 3/4 of (TargetP, Localizer, Multiloc2, PCLR) | 0.622 | 0.791 | 0.647 | 0.905 |
| 4 | 127 | 3/3 of (TargetP, Localizer, PCLR) | 0.588 | 0.822 | 0.644 | 0.906 |
| 5 | 152 | 2/3 of (PredSL, Localizer, Multiloc2) | 0.597 | 0.803 | 0.639 | 0.904 |
| 6 | 68 | 3/4 of (TargetP, PredSL, Localizer, Multiloc2) | 0.575 | 0.827 | 0.639 | 0.905 |
| 7 | 161 | 2/3 of (Localizer, Multiloc2, PCLR) | 0.660 | 0.732 | 0.635 | 0.898 |
| 8 | 189 | 2/2 of (Localizer, PCLR) | 0.634 | 0.756 | 0.634 | 0.900 |
| 9 | 188 | 1/2 of (Localizer, Multiloc2) | 0.697 | 0.696 | 0.632 | 0.894 |
| 10 | 4 | 1 of (Localizer) | 0.675 | 0.714 | 0.632 | 0.896 |
| 11 | 19 | 4/5 of (TargetP, PredSL, Localizer, Multiloc2, PCLR) | 0.563 | 0.828 | 0.632 | 0.903 |
| 12 | 100 | 3/4 of (PredSL, Localizer, Multiloc2, PCLR) | 0.578 | 0.807 | 0.630 | 0.902 |
| 13 | 20 | 3/5 of (TargetP, PredSL, Localizer, Multiloc2, PCLR) | 0.660 | 0.688 | 0.606 | 0.888 |
| 14 | 72 | 3/4 of (TargetP, PredSL, Localizer, PCLR) | 0.648 | 0.697 | 0.606 | 0.889 |
| 15 | 69 | 2/4 of (TargetP, PredSL, Localizer, Multiloc2) | 0.678 | 0.656 | 0.595 | 0.882 |
| 16 | 187 | 2/2 of (Localizer, Multiloc2) | 0.474 | 0.870 | 0.594 | 0.896 |
| 17 | 116 | 2/3 of (TargetP, PredSL, Localizer) | 0.663 | 0.664 | 0.592 | 0.883 |
| 18 | 181 | 2/2 of (PredSL, Localizer) | 0.511 | 0.814 | 0.591 | 0.894 |
| 19 | 160 | 3/3 of (Localizer, Multiloc2, PCLR) | 0.462 | 0.880 | 0.590 | 0.895 |
| 20 | 115 | 3/3 of (TargetP, PredSL, Localizer) | 0.491 | 0.835 | 0.588 | 0.894 |
| 21 | 155 | 2/3 of (PredSL, Localizer, PCLR) | 0.698 | 0.629 | 0.587 | 0.875 |
| 22 | 5 | 1 of (Multiloc2) | 0.495 | 0.826 | 0.587 | 0.894 |
| 23 | 101 | 2/4 of (PredSL, Localizer, Multiloc2, PCLR) | 0.706 | 0.621 | 0.585 | 0.873 |
| 24 | 124 | 3/3 of (TargetP, Localizer, Multiloc2) | 0.452 | 0.883 | 0.585 | 0.894 |
| 25 | 79 | 4/4 of (TargetP, Localizer, Multiloc2, PCLR) | 0.445 | 0.895 | 0.585 | 0.894 |

### Predicted Plastid Proteome Correlates with Genome Size

As a demonstration of the utility of the Localizer and TargetP workflow, subcellular prediction was performed for the whole proteomes of fifteen phylogenetically diverse species. Six monocot species, including *Anthurium amnicola*, *Brachypodium distachyon*, *Oryza sativa*, *Panicum virgatum*, *Setaria italica*, and *Sorghum bicolor* and eight eudicots, including *Arabidopsis thaliana*, *Fragaria vesca*, *Glycine max*, *Malus* × *domestica*, *Populus trichocarpa*, *Prunus persica*, *Solanum lycopersicum*, and *Vitis vinifera* were chosen. Additionally, *Amborella trichopoda*, a species which diverged from the rest of the angiosperms prior to the divergernce of monocots and eudicots, was also incorporated into the comparative analysis. Complete information including data version numbers, proteome sizes, and prediction of plastid-targeted proteins by Localizer and TargetP is summarized in Table 5. In Arabidopsis, 2,826 proteins were predicted to be plastid-targeted, representing 8.8% of all protein isoforms. This finding is in agreement with the conservative estimates of the Arabidopsis plastid proteome [2,4,69]. Similar percentages were calculated in other species but varied from a low of 6.4% in tomato to a high of 9.3% in *A. amnicola*. As expected, the absolute number of predicted plastid-targeted genes showed a high correlation with the genome size (R^2^=0.965) (Figure 2). This result suggests that an increase in genome size and gene content yield a similar increase in the total number of plastid-targeted proteins. Over 10,000 of the Arabidopsis sequences have experimentally-determined localization, and comparing predictions for these sequences revealed an apparent sensitivity of 55.6%, specificity of 89.8%, accuracy of 83.6%, and MCC of 0.614. Sensitivity is somewhat low in this estimation due to the use of MS data, which includes many false positives, but the high specificity suggests good prediction accuracy. With the combination of the high correlation with experimentally-validated proteins and the lack of monocot/eudicot bias imparted by Localizer, it is expected that similar levels of accuracy were achieved for the entire set of species analyzed in this study.

**Figure 2:**
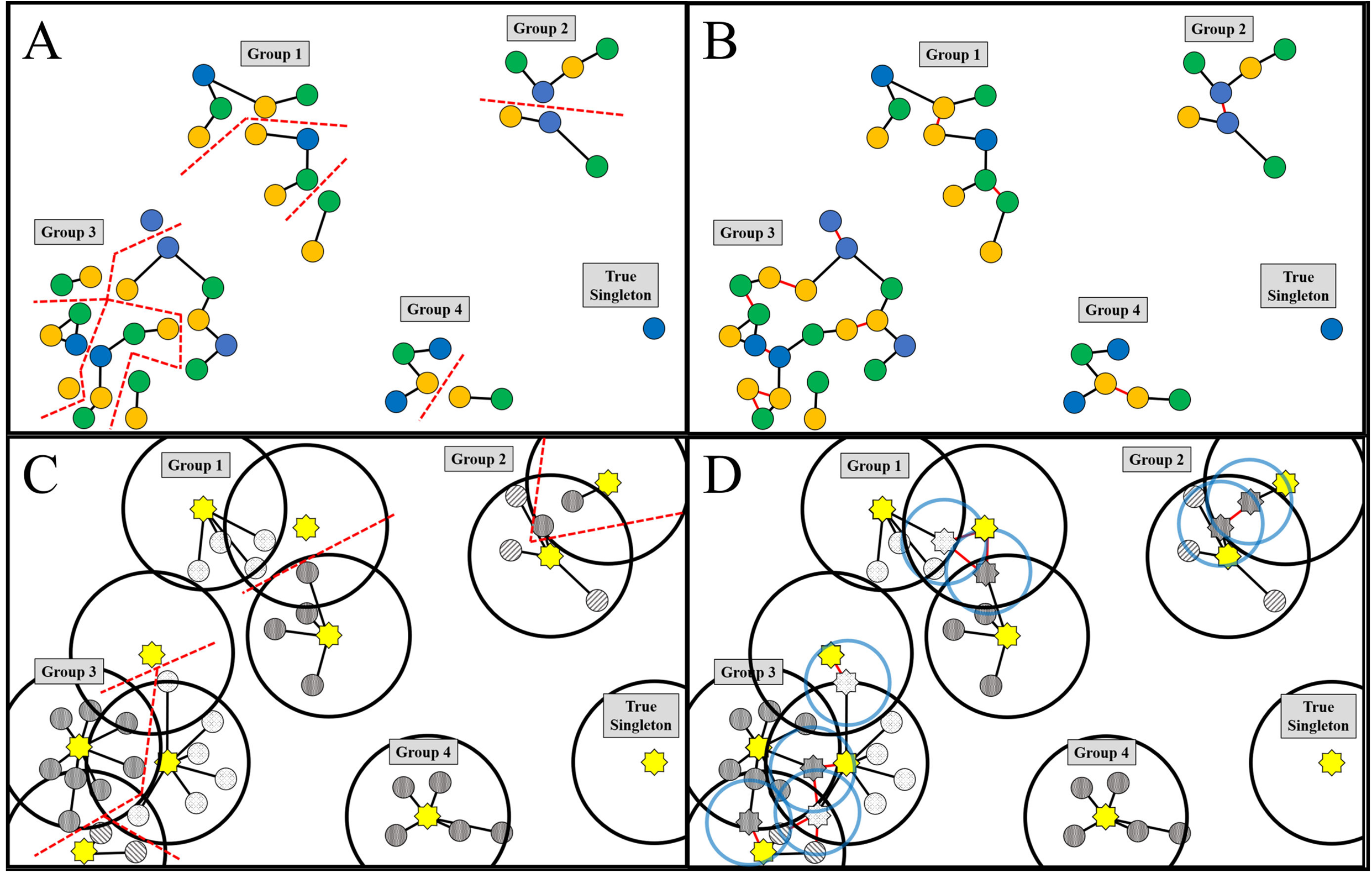
Illustration of RBH and UCLUST Sequence Clustering Methods. Initial (A) and expanded (B) RBH figures indicate clustering between genotypes 1 (blue circles), 2 (green circles), and 3 (orange circles). Bidirectional best BLAST hits between sequences from different genotypes are indicated with black lines; bidirectional better BLAST hits between sequences within the same genotype with red lines and fragments with dotted red lines. For UCLUST, the initial length-sorted (C) run is illustrated with yellow stars indicating centroids, small gray patterned circles indicating non-centroid sequences, large black circles indicating the match range for initial centroids, and black lines indicating sequence clustering for the initial run. For clarity, sequences are patterned to indicate belonging to each initial cluster, and red dotted lines indicate cluster fragmentation. Randomization of centroids (D) mitigates this artificially-induced problem; gray patterned stars indicate randomly-selected centroids, light blue circles indicate the match range for randomly-seeded centroids, and red lines indicate new matches found with red lines. Distances not drawn to scale.

**Table 5:** Targeting Prediction for Selected Genotypes. Predicted protein sequences from fifteen species representing a mixture of model organisms and crop species as well as a mixture of monocots, eudicots, and the early diverging species *Amborella trichopoda* were downloaded from Phytozome (phytozome.jgi.doe.gov) or from the sources indicated in the table. For each genotype, the version, reference, and sequence count are provided from the original publications. *TargetP and Localizer were used to detect plastid-targeted sequences. **Indicates unpublished but publicly-available data downloaded from Phytozome for *Panicum virgatum*.

| Species | Version | Source | Sequences | Plastid-Targeted* |
| --- | --- | --- | --- | --- |
| *Amborella trichopoda* | 1.0 | [70] | 26,846 | 1,833 |
| *Anthurium amnicola* | 1.0 | [71] | 27,959 | 1,324 |
| *Arabidopsis thaliana* | TAIR10 | [72] | 35,386 | 2,826 |
| *Brachypodium distachyon* | 3.1 | [73] | 52,972 | 4,240 |
| *Fragaria vesca* | 1.1 | [74] | 32,831 | 2,051 |
| *Glycine max* | Wm82 | [75] | 73,320 | 5,125 |
| *Malus x domestica* | 1.0<br>(custom transcriptome) | [76]<br>[77–80] | 57,386<br>(74,249) | 4,665 |
| *Oryza sativa* | 7.0 | [81] | 49,061 | 3,417 |
| *Panicum virgatum* | 3.1 | [82]** | 133,775 | 10,262 |
| *Populus trichocarpa* | 3.0 | [83] | 73,013 | 5,741 |
| *Prunus persica* | 2.1 | [84] | 47,089 | 3,615 |
| *Setaria viridis* | 2.2 | [85] | 43,001 | 3,461 |
| *Solanum lycopersicum* | 2.1 | [86] | 47,205 | 1,875 |
| *Sorghum bicolor* | 3.1 | [87] | 34,727 | 3,918 |
| *Vitis vinifera* | 2.0 | [88]<br>[89] | 55,564 | 3,932 |

### Clustering of Gene Families

Although the plastid is highly dependent on proteins imported from the nucleus for normal viability and function, the size and diversity of the plastid proteome across the plant kingdom remain poorly understood. The hypothesis that the plastid proteome is diverse and each species has a unique set of plastid-targeted proteins was examined by grouping sequences into homologous protein groups using two parallel clustering methods (Figure 3). Clustering method has a significant impact on the size and accuracy of the resulting clusters, and therefore on the number and relevance of predictions. Reciprocal best BLAST Hits (RBH) using ALL-v-All BLAST comparisons of whole proteomes are a standard proxy for orthology in comparative genomics, although they are susceptible to inclusion of weakly homologous paralogs. BLAST-based approaches combined with Markov clustering or similar methods to remove paralogs are used in commonly-cited methods such as InParanoid [90], OrthoMCL [91], and COG [92, 93]. However, these methods can bias single-copy genes or highly conserved families which is problematic for polyploid genomes where many-to-many gene relationships are common [94, 95]. For instance, the popular OrthoMCL fails to detect many homologous proteins with conserved expression patterns, and therefore with likely conserved functions, between rice and Arabidopsis [96, 97]. In contrast, more straightforward RBH methods often outperform more complicated algorithms on eukaryotic genomes [98].

**Figure 3:**
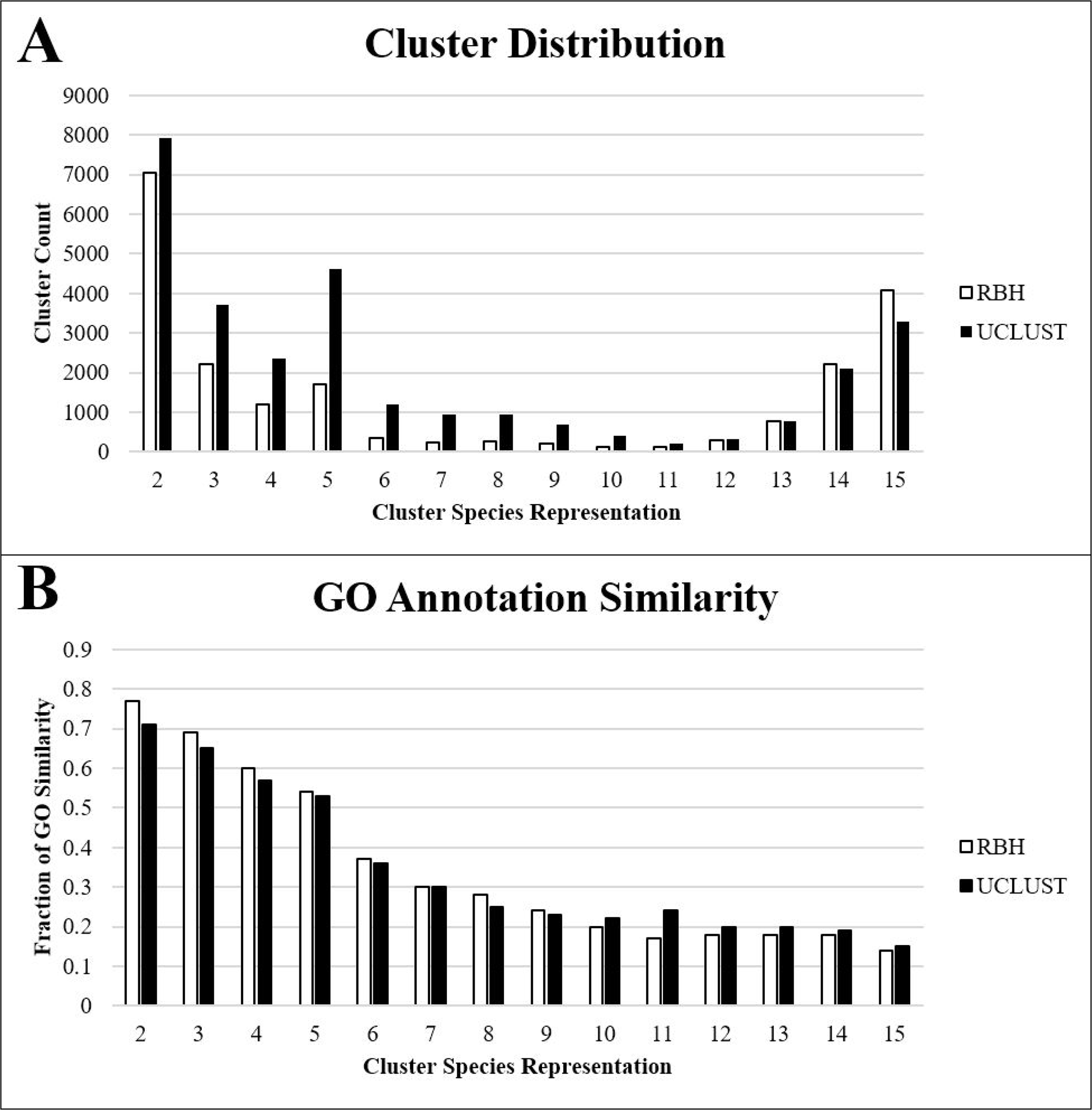
Overall Performance of RBH and UCLUST methods. (A) Cluster distribution in RBH and UCLUST. Both methods resulted in similar distributions of clusters, although RBH resulted in slightly more clusters with 13-15 species and UCLUST resulted in more clusters from 2-12 species. The slight increase in clusters with five species is interesting, and may result from sequences with homology within the Poaceae family or within Rosids but with no significant homologs outside those groups. (B) GO annotation similarity in RBH and UCLUST clusters. Lower similarity scores in higher-order clusters are partially due to different annotation methods and thresholds used for different species. Annotation similarity was generally higher in RBH at smaller cluster sizes and higher in UCLUST for larger clusters. Similarity decreased with the increasing representation of species, which may be partially caused by different annotation methods used for different genome sequencing projects, or may alternatively be caused by decreased homology within large clusters.

A simplified RBH approach, allowing many-to-many relationships, was determined to be most appropriate for this analysis to avoid fracture of gene families with paralogs or co-orthologs. Initial homologous relationships were identified using pairwise BLAST-P comparisons of two genotypes; only sequences which are mutually the best BLAST hits for each other were utilized. Similar methods have used 40% as an appropriate identity and coverage threshold for orthologous relationships [10,99–101]. Therefore 40% was used as the initial threshold of homology. Initial clustering generated many small clusters, so a supplemental method for expansion of clusters, using reciprocal better BLAST hits of each species=’ proteome BLAST’ed against itself, was tested (Additional File 1). A 90% threshold was determined to be optimal for clusters with fewer genotypes decreasing significantly in number, while clusters containing a majority of genotypes remained stable or increased. In contrast, application of between 60 and 80% expansion thresholds caused the liberal merging of clusters into extremely large clusters representing thousands of individual sequences. Additionally, GO term similarity was assessed within clusters at each population size based on the number of species in the cluster and was found to increase slightly for clusters containing few species when using a 90% expansion threshold, while more massive clusters experienced no change or slight decreases.

An alternative approach called UCLUST was implemented to complement the RBH method with a faster and more efficient technique because its semi-global algorithm detects homology in a fraction of the time required for BLAST and becomes much more efficient on enormous datasets. Initial clusters were constructed at a 40% identity and 40% coverage threshold similar to the RBH approach. However, initial clustering produced smaller clusters and resulted in cluster fragmentation. Therefore, modifications were implemented to expand initial clusters by randomly selecting sequences out of each initial cluster and iterating the UCLUST search at more stringent conditions using the selected sequences as new centroids (Additional File 1). Cluster expansion significantly increased the number of clusters with many species, which largely came from the drastic reduction of the number of single-genotype clusters. As with RBH, a 90% expansion threshold was found to be optimal and increased the number of clusters sharing 14-15 species roughly 4-fold, while lower thresholds resulted in the frequent grouping of nonhomologous sequences. Comparison of GO similarity for clusters containing multiple species showed that similarity increased slightly or remained stable for nearly all cluster sizes in the 90% expansion threshold compared to the initial, non-expanded UCLUST analysis. The number of iterations required to fully expand cluster space in UCLUST was also examined, and it was found that most clusters were completely expanded by ten iterations, while further iterations yielded diminishing returns (Additional File 1). A total of 100 iterations were performed to avoid problems with the randomization of centroid sequences.

Application of the optimal clustering methods to the proteomes of the species chosen generated 170,877 clusters using RBH (Table 6) and 103,501 clusters using UCLUST (Table 7). Nearly all the additional clusters in RBH were from single-species clusters or singleton sequences (data not shown): 150,067 of the RBH clusters (87.82%) were single-species clusters of which 134,319 were singleton sequences, while UCLUST detected 74,059 single-species clusters (71.55%) including 45,033 singletons. Some of these may be orphan genes, but they are more likely to be prediction and annotation errors or pseudogenes because the lack of homology implies lack of conserved function or extreme mutation rates that are more likely to occur in non-coding sequences. A total of 20,810 and 29,442 clusters in RBH and UCLUST approach, respectively, contained sequences from multiple species; while they represented a minority of clusters, they contained the majority of initial sequences. A bimodal distribution was observed in both methods in which two clusters, the first containing 14-15 of the species and the second containing just 2-3 species, represented the majority of the clusters (Figure 3A). Comparatively fewer clusters contained between 4-13 species. Of the conserved clusters containing all 15 species, RBH detected 4,090 clusters, while UCLUST yielded 3,295. GO similarity between UCLUST and RBH was remarkably consistent, but UCLUST had somewhat better scores for conserved clusters containing plastid-targeted sequences from all species and lower scores for semi-conserved or non-conserved clusters containing few species (Figure 4B). Across both methods, GO similarity decreased with increasing cluster size. While the merging of nonhomologous sequences may be partially responsible for this decrease, the annotation methods and parameters are not identical for the species used in this study, which artificially decreases the apparent similarity score regardless of clustering specificity.

**Figure 4:**
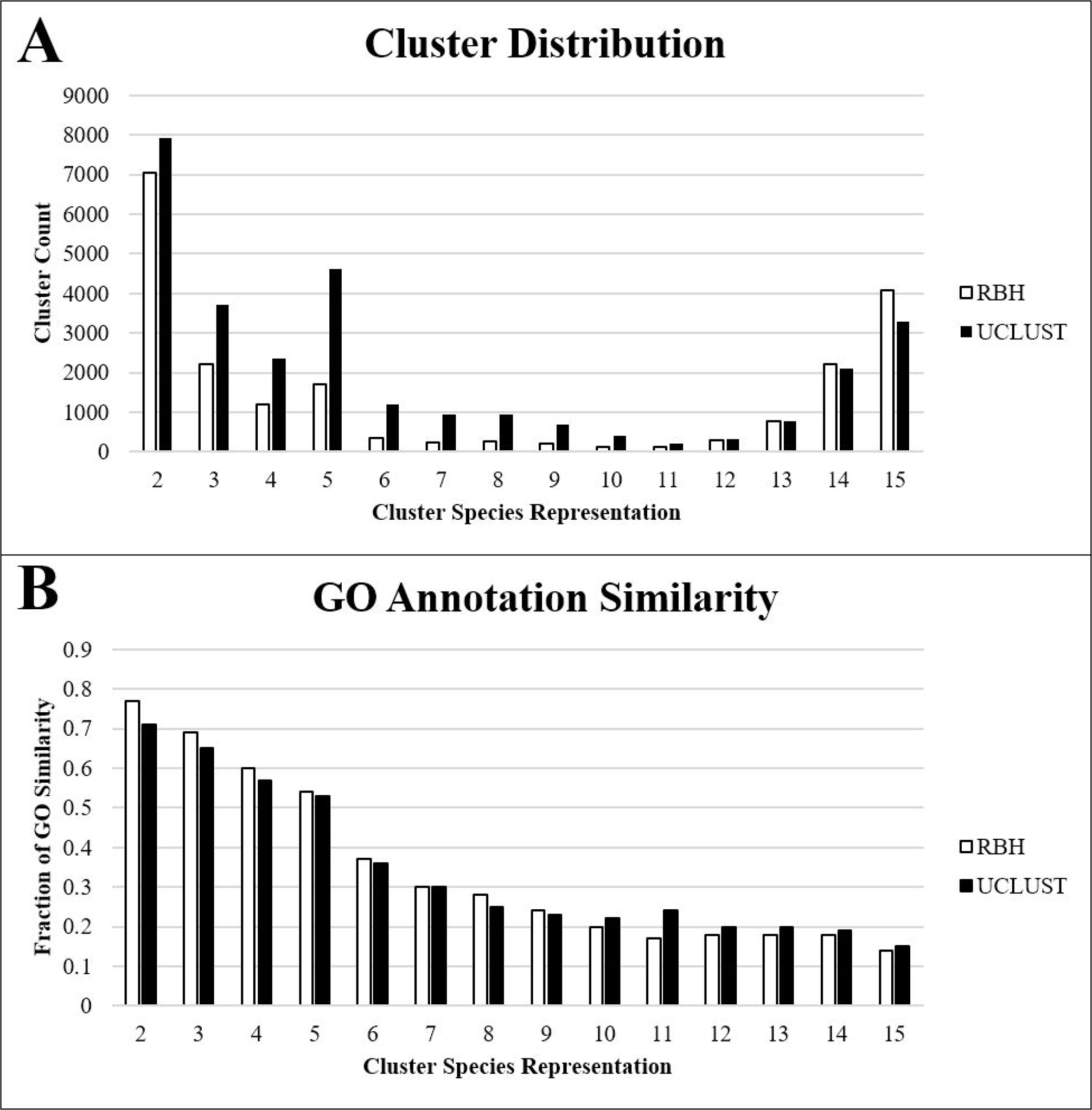
Workflow Diagram of Sequence Clustering Methods. For RBH (left panel), 1. initial cluster edges were generated by finding all reciprocal best-BLAST hits in all-v-all comparisons of proteomes from two separate species at a 40% identity, 40% coverage threshold, and 2. Secondary cluster edges were generated by finding all reciprocal better-BLAST hits in all-v-all comparisons of each proteome against itself at a 90% identity, 90% coverage threshold. For UCLUST (right panel), 1. An initial run was performed at 40% identity and 40% coverage threshold on a FASTA file containing sequences from every species in length-sorted order, and 2. Random sequences of at least 90% identity and 90% coverage were extracted from each cluster, this subset was length-sorted, and then the original length-sorted FASTA file was concatenated to the new seed sequences. This process was iterated 100 times, and a separate UCLUST run was performed for each iteration. Downstream processes for RBH and UCLUST were identical: 3. All clusters/pairs with a shared sequence were condensed into single clusters, 4. All sequences that failed to have at least 40% identity and 40% coverage based on BLAST-P analysis to any of the predicted plastid-targeted sequences in the cluster were trimmed out, 5A. all clusters with at least three species were extracted, and 5B. Clusters containing plastid-targeted sequences were sorted into “conserved,” “semi-conserved,” and “non-conserved” groups according to the number of species with predicted plastid targeting and the taxonomic grouping of those species. cTP – chloroplast transit peptide.

**Table 6:**
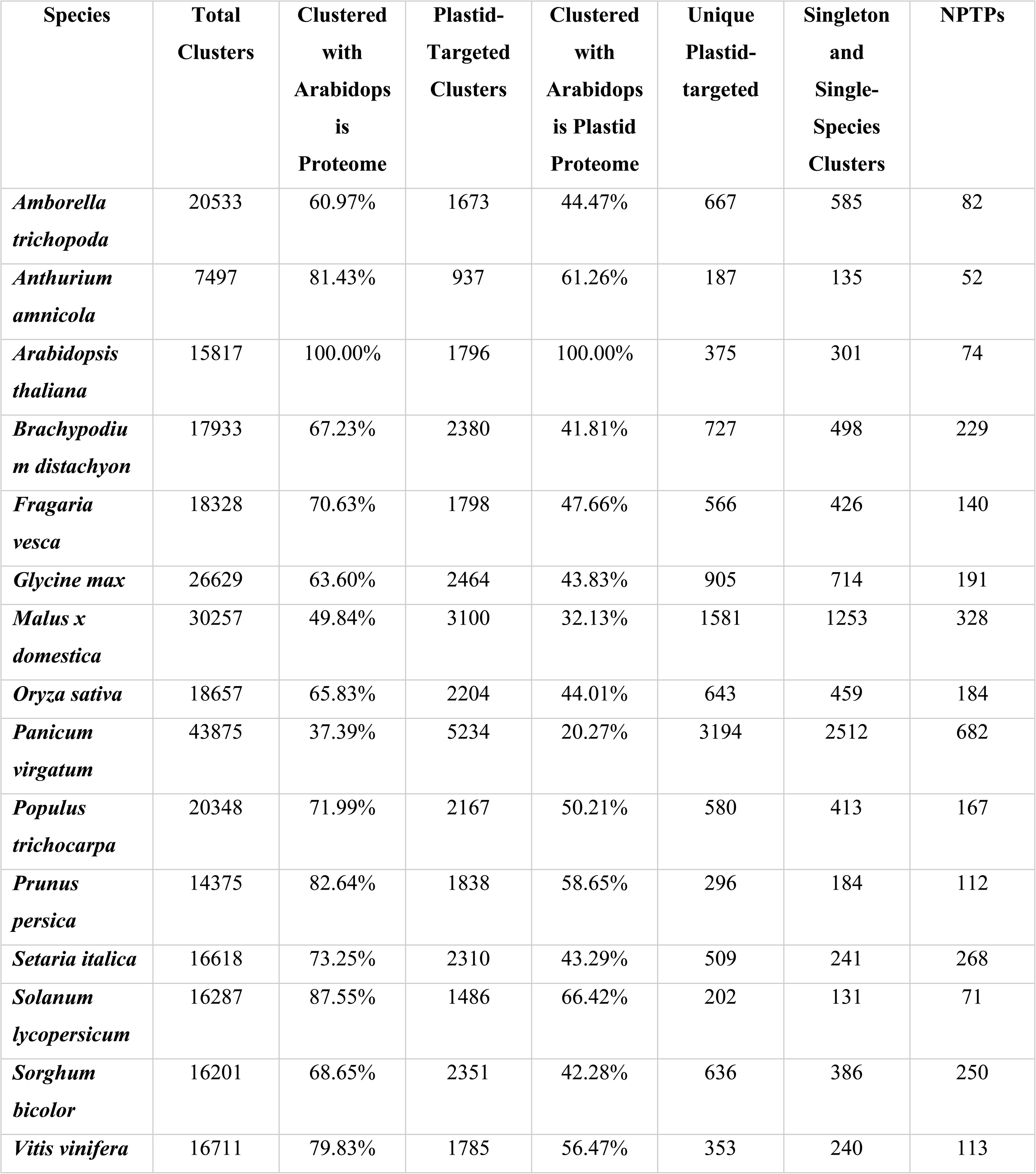
RBH Clustering Results by Genotype. Clustering of gene families using 40% reciprocal Intergeneric best BLAST hits and 90% reciprocal Intergeneric better BLAST hits was performed, and clusters containing plastid-targeted sequences were identified for each genotype. The number of total proteomes and plastid-targeted clusters with at least one Arabidopsis sequence were identified, as well as the number of clusters containing a plastid-targeted sequence from only the selected genotype. The number clusters overlapping with Arabidopsis for all clusters and plastid-targeted clusters was identified, as well as the number of clusters containing a plastid-targeted sequence from only the selected genotype. NPTPs – Nascent Plastid Targeted Proteins.

**Table 7:** UCLUST Clustering Results by Genotype. Clustering of gene families was performed using an initial UCLUST iteration with 40% coverage and 40% identity followed by extraction of random sequences from each cluster to seed additional iterations performed at 90% coverage and identity. Clusters containing shared sequences were merged, followed by identification of clusters containing plastid-targeted sequences in each genotype. The number clusters overlapping with Arabidopsis for all clusters and plastid-targeted clusters was identified, as well as the number of clusters containing a plastid-targeted sequence from only the selected genotype.

| Species | Total Clusters | Cluster ed with Arabido psis Proteo me | Plastid-Targeted Clusters | Clustered with Arabidopsis Plastid proteome | Unique Plastid-targeted | Singleton and Single-Species Clusters | NPT Ps |
| --- | --- | --- | --- | --- | --- | --- | --- |
| *Amborella trichopoda* | 19190 | 55.61% | 1721 | 41.78% | 736 | 541 | 195 |
| *Anthurium amnicola* | 7365 | 78.22% | 909 | 58.53% | 173 | 76 | 97 |
| *Arabidopsis thaliana* | 13065 | 100.00 % | 1783 | 100.00% | 261 | 95 | 166 |
| *Brachypodium distachyon* | 16777 | 57.05% | 2375 | 37.68% | 623 | 225 | 398 |
| *Fragaria vesca* | 16821 | 65.75% | 1828 | 46.01% | 551 | 172 | 379 |
| *Glycine max* | 20157 | 70.78% | 2320 | 49.31% | 637 | 296 | 341 |
| *Malus x domestica* | 21427 | 56.25% | 2846 | 35.45% | 1197 | 469 | 728 |
| *Oryza sativa* | 18102 | 55.76% | 2249 | 38.86% | 564 | 235 | 329 |
| *Panicum virgatum* | 29207 | 34.12% | 4725 | 20.00% | 2506 | 1048 | 1458 |
| *Populus trichocarpa* | 15881 | 78.35% | 1977 | 56.35% | 335 | 95 | 240 |
| *Prunus persica* | 14753 | 79.49% | 1921 | 57.16% | 229 | 36 | 193 |
| *Setaria italica* | 17810 | 57.11% | 2427 | 36.51% | 480 | 98 | 382 |
| *Solanum lycopersicum* | 15675 | 81.13% | 1574 | 62.26% | 245 | 79 | 166 |
| *Sorghum bicolor* | 17395 | 54.56% | 2410 | 36.14% | 554 | 191 | 363 |
| *Vitis vinifera* | 16092 | 79.06% | 1805 | 57.62% | 299 | 102 | 197 |

### Identification of Gene Families with Conserved Plastid Targeting

Genomes of endosymbiotic bacteria contain 1,500 proteins on average, and plastids are likely to contain similar numbers when accounting for both the plastid genome and core nuclear-encoded plastid-targeted genes [102]. To determine the number of gene families with conserved plastid localization, clusters containing at least 13 species, of which all species contained at least one predicted plastid-targeted sequence or at least four non-plastid-targeted sequences were selected. These parameters were chosen to account for assembly and annotation errors and to correct for the 39% false negative prediction rate for bonafide plastid-targeted proteins which could eliminate many truly conserved clusters. There is a nearly 20% chance that at least one of four random sequences with non-plastid localization prediction is a false negative, but sequences that already share homology to predicted plastid-targeted sequences have a significantly higher likelihood of being false negatives. A workflow diagram representing cluster detection, filtering, processing, and categorization is represented in Figure 4. Applying this workflow, 628 conserved protein clusters were found in RBH (Table 6, Figure 5), while UCLUST detected 828 (Table 7, Figure 6). Of these, 621 clusters in RBH and 817 in UCLUST also contain sequences from *A. trichopoda*, and all have several monocot and eudicot sequences, strongly indicating that these clusters represent the fundamental core plastid-targeted gene families. Previous estimates predicted that 857-1020 sequences were shared between rice and Arabidopsis, another report projected that between 289-737 proteins were shared among the chloroplast proteomes of seven plant species [2, 10]. Identification of gene families with conserved chloroplast transit peptides is an essential output of this work, as *in silico* methods can quickly identify conserved plastid-targeted proteins that have failed to be detected by genetic screens due to embryo lethality, gene redundancy, or random chance. Several methods have validated these sequences as truly plastid-targeted and representative of conserved plastid-targeted genes. First, Arabidopsis proteins with experimentally-validated localization were examined within the conserved clusters. A total of 84.2% (183 proteins) of predicted plastid-targeted Arabidopsis sequences in conserved RBH clusters were validated by GFP and 94.5% (1,054) were validated by MS. The same was true for 80.5% (154 proteins) and 92% (855 proteins) in conserved RBH and UCLUST clusters, respectively (Additional Files 4 and 5). While these methods have yielded good overall sensitivity, small errors at initial stages of clustering can compound in larger clusters and result in unrealistically high numbers of sequences. For RBH, an average of 113.9 sequences and median of 61 were present in conserved clusters while UCLUST produced an average of 125.9 sequences and median of 84. Most sequences in these clusters come from a small set of genotypes: *G. max*, *P. virgatum*, *P. trichocarpa*, and *V. vinifera* each contributed an average of over 10 sequences each to clusters with shared plastid localization prediction, while *M.* × *domestica* contributed over 10 sequences on average in UCLUST (summarized in Additional Files 4 and 5). Significant gene duplication or inclusion of multiple gene isoforms especially in those species likely accounts for a portion of the larger cluster sizes, but more distantly paralogous sequences which are less likely to share biological function are also likely to be common. Thus, the list of conserved clusters reported here is not meant to be definitive and final, but rather a general guide which will require phylogenetic and experimental validation. In cases where larger clusters contain multiple paralogs or non-homologous, phylogenetic methods could resolve homology relationship with higher efficiency than the currently used RBH and UCLUST methods. However, the biological accuracy of the predicted plastid-targeted sequences within these clusters is still high.

**Figure 5:**
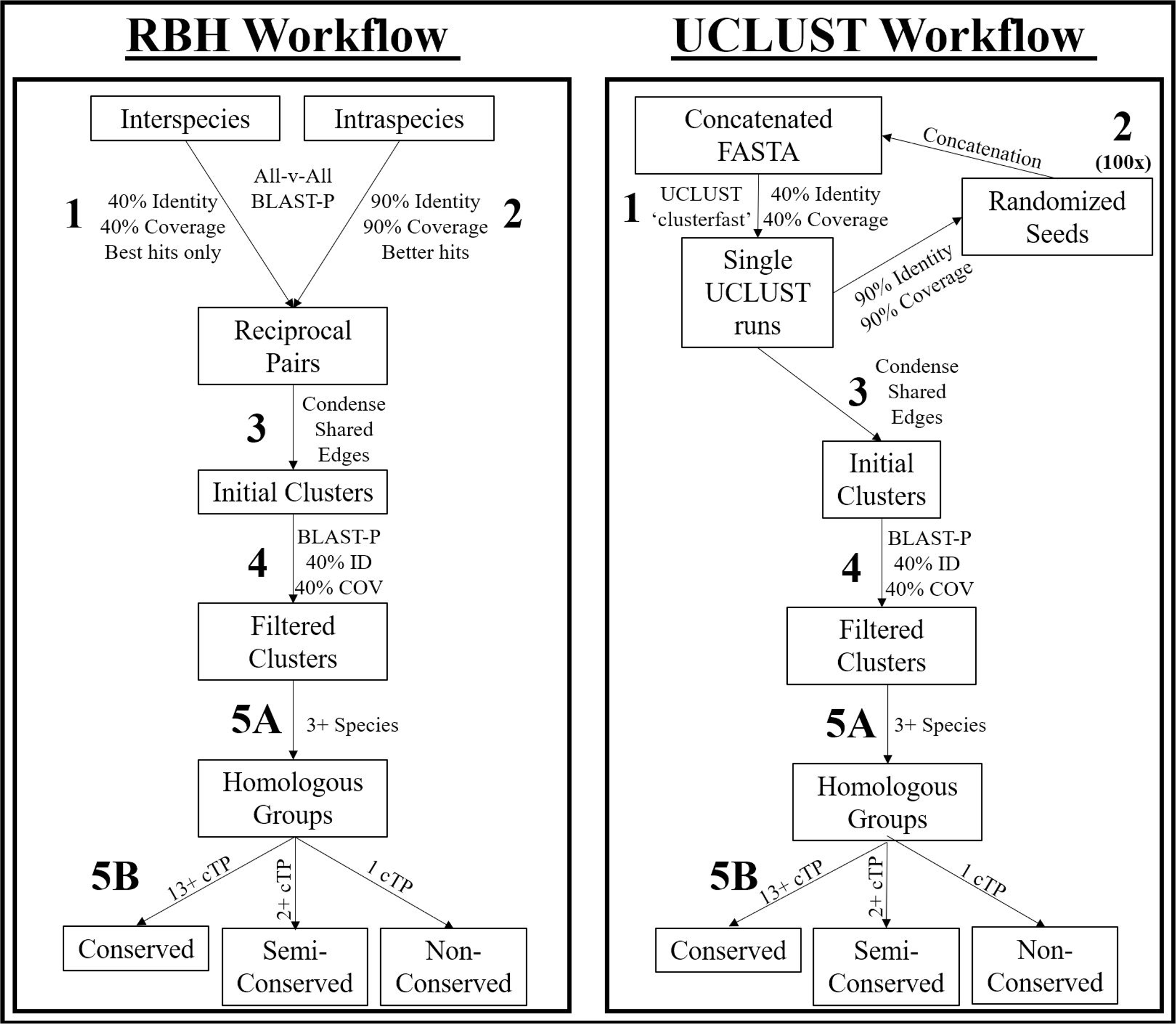
RBH Visual Representation. For “unique” clusters, single-species and singleton clusters are not represented, leaving only clusters with non-targeted homologs present in other species. The relative size of these unique clusters is represented by the area of the respective circle. Shared protein groups at the kingdom, clade, subclade, and family levels are not represented by figure size. Overall, 628 protein clusters were shared between all 15 species, 1,002 had plastid-targeting specific to either monocots or eudicots, and 2,943 had plastid-targeting specific to only a single species.

**Figure 6:**
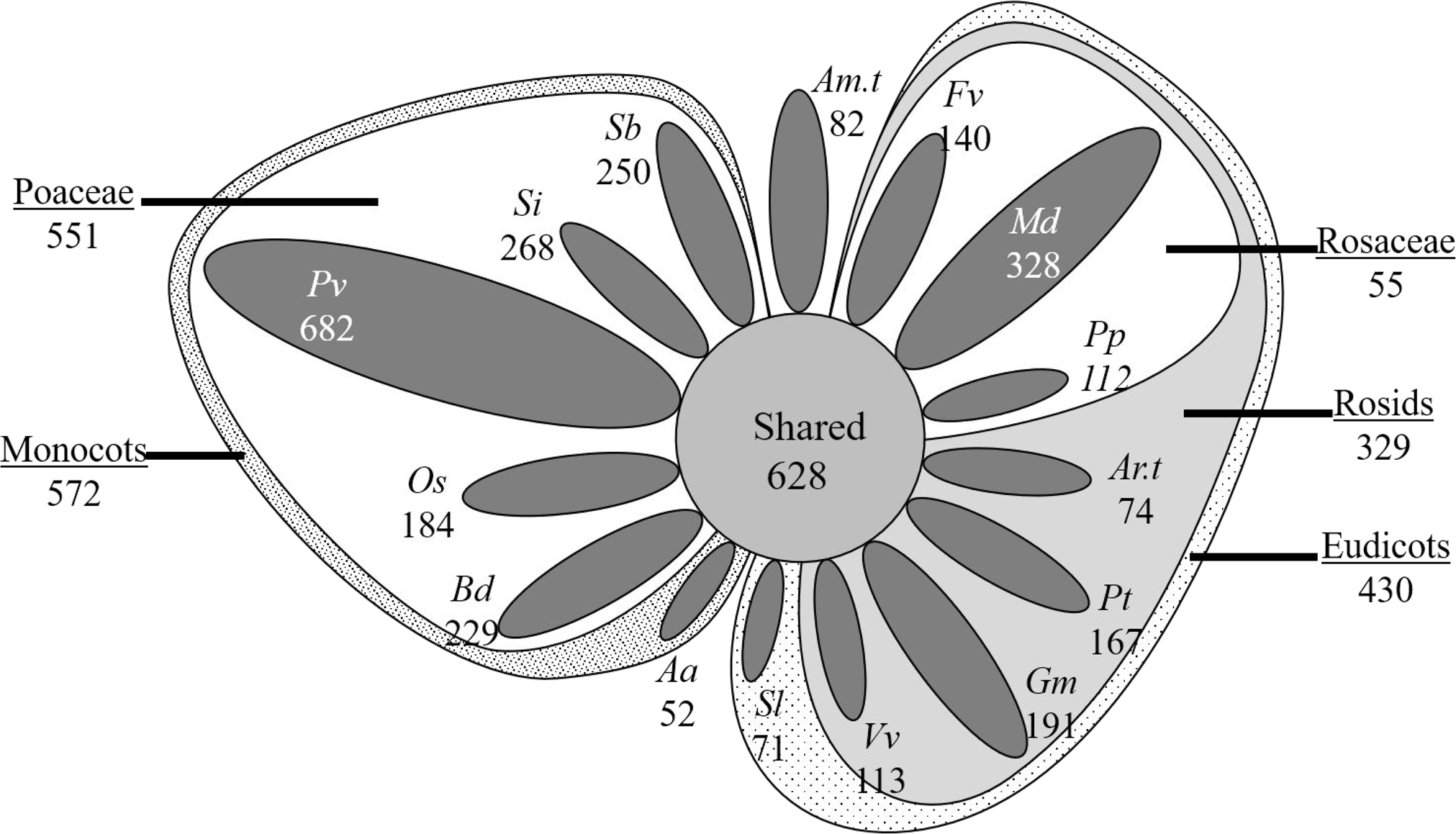
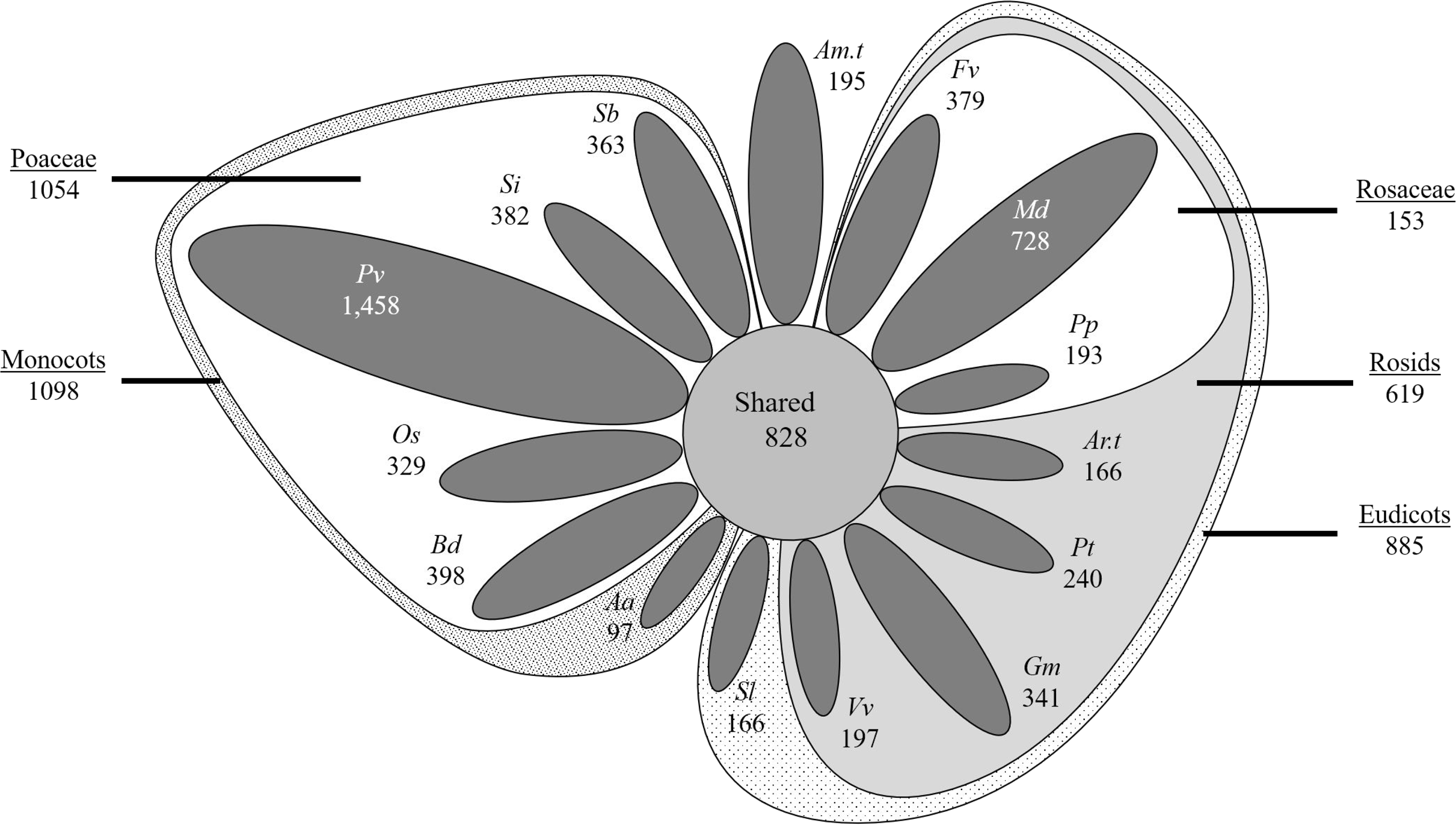
UCLUST Visual Representation. For “unique” clusters, single-species and singleton clusters are not represented, leaving only clusters with non-targeted homologs present in other species. The relative size of these unique clusters is represented by the area of the respective circle. Shared protein groups at the kingdom, clade, subclade, and family levels are not represented by figure size. Overall, 828 protein clusters included plastid-targeted sequences from all 15 species, 1,983 had plastid-targeting specific to monocots or eudicots, and 5,632 had plastid-targeting specific to a single species.

Next, enrichment of gene ontology (GO) annotations was performed in conserved clusters by finding GO terms shared in at least three individual sequences and for over 10% of sequences. Terms were compared to annotations extracted using the same criteria for all the clusters of the respective clustering method and GO term enrichment was performed using BLAST2GO [103]. Overall, 53 terms including 29 terms associated with biological function, 23 associated with the cellular component, and one associated with the molecular process were found for RBH (Table 8). In UCLUST, a total of 33 terms were found, including 15 associated with the biological process, 17 with the cellular component, and one with the molecular process (Table 9). The most significantly enriched GO terms under the biological process ontology for both RBH and UCLUST methods were GO:0015979 (photosynthesis) and GO:0008152 (metabolic process), while a majority of the remaining highly enriched terms were associated with homeostatic processes (GO:0042592), cellular component organization (GO:0016043), single-organism biosynthetic processes (GO:0016043), generation of precursor metabolites (GO:0006091), and lipid metabolism (GO:0006629). In the RBH method, additional terms associated with amide, peptide, and organonitrogen compound biosynthesis and metabolism (GO:0043604, GO:0043603, GO:0043043, GO:0006518, GO:1901566, GO:1901564, GO:0044271, GO:0034641, GO:0006807), were enriched. UCLUST additionally had enriched GO terms associated with transport (GO:0006810), localization (GO:0051234, GO:0051179) and metabolism of carbohydrates (GO:0005975). Among cellular component ontologies, plastid (GO:0009536) was the most overrepresented term in both methods. Other highly overrepresented cellular component terms included organelle (GO:0043226), thylakoid (GO:0009579), chloroplast (GO:0009507), and associated terms. In RBH methods, significant enrichment of ribonucleoprotein complexes (GO:1990904, GO:0030529) was found. For the molecular process ontology, structural molecule activity (GO:0005198) was enriched in RBH and catalytic activity (GO:0003824) in UCLUST. These GO terms were further compared to the results of a previous study involving intergeneric analysis that described 737 conserved plastid-targeted proteins [10]. In this study, 42% of enriched terms found using UCLUST overlapped with the methods reported previously [10]. RBH methods were somewhat lower because more enriched terms were found, but still overlapped with the previously published dataset by 24%. These results are remarkably similar given that only GO terms from Arabidopsis had been examined previously and also different methods of GO enrichment had been used in those studies. The final and perhaps the most important test of the biological significance of conserved plastid-targeted clusters is whether they contain proteins expected to be present in plastids of all higher plants. Gene names were retrieved from TAIR10 for all Arabidopsis sequences in conserved clusters, and many of the most prominent plastid proteins were confirmed to be present in clusters for both RBH and UCLUST methods. The following is not intended to be an exhaustive list but merely a representative of the types of proteins detected in conserved plastid-targeted clusters; a complete list of annotations and gene names in RBH and UCLUST clusters are available in Additional Files 4 and 5. Among genes involved in primary photosynthesis, HCEF, LhcA1, LhcA2, LhcB1, LhcB2, LhcB3, Lhcb4, LPA1, LPA3, PPDK, and RbcS were detected in both methods, while LPA66 was found in RBH only. Photosystem subunits Psa-E, Psa-F, Psa-G, Psa-H, Psa-K, Psa-N, PsbP, Psa-O, PsbQ, PsbR, PsbS, PsbW, and PsbY were also found in both methods, while PsbT-N and PsbX were found only in RBH and PsbO was found only in UCLUST. Among ribosomal proteins, Rps1, Rps9, Rpl4, Rpl11, and Rpl12 were detected by both techniques, while Rpl9 and Rpl15 were only found using RBH and Rpl10 was found only with UCLUST. Proteins involved in translocation and chaperone functions found by both methods included ClpB, ClpC, ClpD, ClpP, ClpR, FtsH, Hsp60, Hsp70, Hsp88, Hsp90, Hsp98, Cpn10, Cpn20, Cpn60, Vipp1, Alb3, Alb4, TatC, Tic20, Tic21, Tic40, Tic55, Tic110, Toc75, and Plsp1. The Sec translocase subunits SecA, Scy1, and Scy2 were uniquely found in RBH, while organellar oligopeptidase OOP was also found in UCLUST. Finally, genes associated with primary plastid metabolism (SBPase, TPT, FRUCT5, G6PD2, and G6PD), heme biosynthesis (GUN2, GUN5, HEMA, HEMB, HEMC, HY2, PORA, PORB, and PORC), and fatty acid synthesis (ACC2, FAB2, FAD7, FAD8, FATA, FATB, lipoxygenase) were found in core clusters.

**Table 8:** Enriched GO terms for Conserved Plastid-Targeted RBH Clusters. . Clusters containing at least 13 species with predicted or likely plastid-targeted sequences were mined for common GO terms and compared against terms extracted for the total set of RBH-derived clusters using BLAST2GO. All terms enriched above p=1.0E^-5^ in core plastid-targeted clusters are presented below.

|  | GO term | Description | Ontology | P-value | FDR |
| --- | --- | --- | --- | --- | --- |
| 1 | GO:0015979 | photosynthesis | BIOLOGICAL_PROCESS | 1.73E-44 | 3.99E-47 |
| 2 | GO:0008152 | metabolic process | BIOLOGICAL_PROCESS | 6.40E-27 | 1.59E-29 |
| 3 | GO:0006091 | generation of precursor metabolites and energy | BIOLOGICAL_PROCESS | 1.43E-24 | 3.82E-27 |
| 4 | GO:0009058 | biosynthetic process | BIOLOGICAL_PROCESS | 3.73E-20 | 1.20E-22 |
| 5 | GO:0044711 | single-organism biosynthetic process | BIOLOGICAL_PROCESS | 3.00E-15 | 1.02E-17 |
| 6 | GO:0016043 | cellular component organization | BIOLOGICAL_PROCESS | 4.28E-12 | 1.64E-14 |
| 7 | GO:0071840 | cellular component organization or biogenesis | BIOLOGICAL_PROCESS | 6.51E-12 | 2.66E-14 |
| 8 | GO:0044710 | single-organism metabolic process | BIOLOGICAL_PROCESS | 1.05E-10 | 5.06E-13 |
| 9 | GO:0006629 | lipid metabolic process | BIOLOGICAL_PROCESS | 3.16E-10 | 1.57E-12 |
| 10 | GO:0043604 | amide biosynthetic process | BIOLOGICAL_PROCESS | 7.84E-09 | 4.05E-11 |
| 11 | GO:0043603 | cellular amide metabolic process | BIOLOGICAL_PROCESS | 9.01E-09 | 4.81E-11 |
| 12 | GO:0019725 | cellular homeostasis | BIOLOGICAL_PROCESS | 1.07E-08 | 5.93E-11 |
| 13 | GO:0044699 | single-organism process | BIOLOGICAL_PROCESS | 1.22E-08 | 7.39E-11 |
| 14 | GO:0009987 | cellular process | BIOLOGICAL_PROCESS | 1.22E-08 | 7.21E-11 |
| 15 | GO:0065008 | regulation of biological quality | BIOLOGICAL_PROCESS | 1.54E-08 | 9.56E-11 |
| 16 | GO:0006412 | translation | BIOLOGICAL_PROCESS | 1.67E-08 | 1.10E-10 |
| 17 | GO:0042592 | homeostatic process | BIOLOGICAL_PROCESS | 1.67E-08 | 1.08E-10 |
| 18 | GO:0043043 | peptide biosynthetic process | BIOLOGICAL_PROCESS | 1.73E-08 | 1.17E-10 |
| 19 | GO:0006518 | peptide metabolic process | BIOLOGICAL_PROCESS | 1.80E-08 | 1.25E-10 |
| 20 | GO:1901566 | organonitrogen compound biosynthetic process | BIOLOGICAL_PROCESS | 2.95E-08 | 2.10E-10 |
| 21 | GO:1901564 | organonitrogen compound metabolic process | BIOLOGICAL_PROCESS | 1.04E-07 | 7.58E-10 |
| 22 | GO:0034641 | cellular nitrogen compound metabolic process | BIOLOGICAL_PROCESS | 1.61E-05 | 1.52E-07 |
| 23 | GO:0044271 | cellular nitrogen compound biosynthetic process | BIOLOGICAL_PROCESS | 1.93E-05 | 1.85E-07 |
| 24 | GO:0006807 | nitrogen compound metabolic process | BIOLOGICAL_PROCESS | 2.26E-05 | 2.21E-07 |
| 25 | GO:0044249 | cellular biosynthetic process | BIOLOGICAL_PROCESS | 2.52E-05 | 2.51E-07 |
| 26 | GO:1901576 | organic substance biosynthetic process | BIOLOGICAL_PROCESS | 4.41E-05 | 4.55E-07 |
| 27 | GO:0034645 | cellular macromolecule biosynthetic process | BIOLOGICAL_PROCESS | 4.47E-05 | 4.69E-07 |
| 28 | GO:0009059 | macromolecule biosynthetic process | BIOLOGICAL_PROCESS | 4.74E-05 | 5.06E-07 |
| 29 | GO:0010467 | gene expression | BIOLOGICAL_PROCESS | 2.96E-04 | 3.31E-06 |
| 30 | GO:0009536 | plastid | CELLULAR_COMPONENT | 1.35E-279 | 2.41E-283 |
| 31 | GO:0005622 | intracellular | CELLULAR_COMPONENT | 1.07E-222 | 3.82E-226 |
| 32 | GO:0044424 | intracellular part | CELLULAR_COMPONENT | 1.22E-222 | 6.49E-226 |
| 33 | GO:0044464 | cell part | CELLULAR_COMPONENT | 6.98E-222 | 6.21E-225 |
| 34 | GO:0005623 | cell | CELLULAR_COMPONENT | 6.98E-222 | 5.62E-225 |
| 35 | GO:0005737 | cytoplasm | CELLULAR_COMPONENT | 2.03E-218 | 2.16E-221 |
| 36 | GO:0044444 | cytoplasmic part | CELLULAR_COMPONENT | 1.38E-217 | 1.72E-220 |
| 37 | GO:0043229 | intracellular organelle | CELLULAR_COMPONENT | 1.94E-193 | 2.76E-196 |
| 38 | GO:0043226 | organelle | CELLULAR_COMPONENT | 1.97E-193 | 3.16E-196 |
| 39 | GO:0043231 | intracellular membrane-bounded organelle | CELLULAR_COMPONENT | 1.16E-179 | 2.06E-182 |
| 40 | GO:0043227 | membrane-bounded organelle | CELLULAR_COMPONENT | 5.74E-179 | 1.12E-181 |
| 41 | GO:0009579 | thylakoid | CELLULAR_COMPONENT | 2.71E-68 | 5.78E-71 |
| 42 | GO:0016020 | membrane | CELLULAR_COMPONENT | 1.08E-20 | 3.06E-23 |
| 43 | GO:0005739 | mitochondrion | CELLULAR_COMPONENT | 2.48E-12 | 8.82E-15 |
| 44 | GO:0005840 | ribosome | CELLULAR_COMPONENT | 1.48E-11 | 6.31E-14 |
| 45 | GO:1990904 | ribonucleoprotein complex | CELLULAR_COMPONENT | 3.36E-11 | 1.55E-13 |
| 46 | GO:0030529 | intracellular ribonucleoprotein complex | CELLULAR_COMPONENT | 3.36E-11 | 1.55E-13 |
| 47 | GO:0032991 | macromolecular complex | CELLULAR_COMPONENT | 1.22E-08 | 7.29E-11 |
| 48 | GO:0009507 | chloroplast | CELLULAR_COMPONENT | 2.01E-07 | 1.61E-09 |
| 49 | GO:0043228 | non-membrane-bounded organelle | CELLULAR_COMPONENT | 3.31E-06 | 3.06E-08 |
| 50 | GO:0043232 | intracellular non-membrane-bounded organelle | CELLULAR_COMPONENT | 3.31E-06 | 3.06E-08 |
| 51 | GO:0044434 | chloroplast part | CELLULAR_COMPONENT | 3.09E-04 | 3.52E-06 |
| 52 | GO:0044435 | plastid part | CELLULAR_COMPONENT | 3.34E-04 | 3.86E-06 |
| 53 | GO:0005198 | structural molecule activity | MOLECULAR_FUNCTION | 3.04E-05 | 3.08E-07 |

**Table 9:** Enriched GO terms for Conserved Plastid-Targeted UCLUST Clusters. . Clusters containing at least 13 species with predicted or likely plastid-targeted sequences were mined for common GO terms and compared against terms extracted for the total set of UCLUST -derived clusters using BLAST2GO. All terms enriched above p=1.0E^-5^ in core plastid-targeted clusters are presented below.

|  | GO term | Description | Ontology | P-value | FDR |
| --- | --- | --- | --- | --- | --- |
| 1 | GO:0008152 | metabolic process | BIOLOGICAL_PROCESS | 3.36E-32 | 9.19E-35 |
| 2 | GO:0015979 | photosynthesis | BIOLOGICAL_PROCESS | 1.24E-29 | 3.62E-32 |
| 3 | GO:0044710 | single-organism metabolic process | BIOLOGICAL_PROCESS | 1.38E-21 | 4.57E-24 |
| 4 | GO:0044711 | single-organism biosynthetic process | BIOLOGICAL_PROCESS | 5.52E-16 | 2.15E-18 |
| 5 | GO:0044699 | single-organism process | BIOLOGICAL_PROCESS | 2.89E-15 | 1.18E-17 |
| 6 | GO:0006091 | generation of precursor metabolites and energy | BIOLOGICAL_PROCESS | 1.15E-13 | 4.93E-16 |
| 7 | GO:0005975 | carbohydrate metabolic process | BIOLOGICAL_PROCESS | 1.20E-10 | 5.84E-13 |
| 8 | GO:0006629 | lipid metabolic process | BIOLOGICAL_PROCESS | 1.33E-06 | 7.01E-09 |
| 9 | GO:0051234 | establishment of localization | BIOLOGICAL_PROCESS | 1.85E-04 | 1.23E-06 |
| 10 | GO:0006810 | transport | BIOLOGICAL_PROCESS | 1.85E-04 | 1.20E-06 |
| 11 | GO:0051179 | localization | BIOLOGICAL_PROCESS | 2.65E-04 | 1.81E-06 |
| 12 | GO:0016043 | cellular component organization | BIOLOGICAL_PROCESS | 2.98E-04 | 2.10E-06 |
| 13 | GO:0044723 | single-organism carbohydrate metabolic process | BIOLOGICAL_PROCESS | 3.01E-04 | 2.23E-06 |
| 14 | GO:0071840 | cellular component organization or biogenesis | BIOLOGICAL_PROCESS | 4.29E-04 | 3.26E-06 |
| 15 | GO:0042592 | homeostatic process | BIOLOGICAL_PROCESS | 8.59E-04 | 6.87E-06 |
| 16 | GO:0009536 | plastid | CELLULAR_COMPONENT | 1.01E-165 | 1.97E-169 |
| 17 | GO:0044464 | cell part | CELLULAR_COMPONENT | 1.54E-140 | 6.02E-144 |
| 18 | GO:0005623 | cell | CELLULAR_COMPONENT | 1.52E-139 | 8.87E-143 |
| 19 | GO:0044444 | cytoplasmic part | CELLULAR_COMPONENT | 8.76E-120 | 6.83E-123 |
| 20 | GO:0005737 | cytoplasm | CELLULAR_COMPONENT | 3.57E-119 | 3.49E-122 |
| 21 | GO:0044424 | intracellular part | CELLULAR_COMPONENT | 2.06E-110 | 2.41E-113 |
| 22 | GO:0005622 | intracellular | CELLULAR_COMPONENT | 1.27E-104 | 1.73E-107 |
| 23 | GO:0043229 | intracellular organelle | CELLULAR_COMPONENT | 6.39E-93 | 1.05E-95 |
| 24 | GO:0043226 | organelle | CELLULAR_COMPONENT | 6.39E-93 | 1.12E-95 |
| 25 | GO:0043231 | intracellular membrane-bounded organelle | CELLULAR_COMPONENT | 6.87E-82 | 1.34E-84 |
| 26 | GO:0043227 | membrane-bounded organelle | CELLULAR_COMPONENT | 1.90E-81 | 4.07E-84 |
| 27 | GO:0009579 | thylakoid | CELLULAR_COMPONENT | 4.05E-39 | 9.47E-42 |
| 28 | GO:0016020 | membrane | CELLULAR_COMPONENT | 4.66E-36 | 1.18E-38 |
| 29 | GO:0071944 | cell periphery | CELLULAR_COMPONENT | 7.58E-11 | 3.55E-13 |
| 30 | GO:0005886 | plasma membrane | CELLULAR_COMPONENT | 1.32E-07 | 6.69E-10 |
| 31 | GO:0009507 | chloroplast | CELLULAR_COMPONENT | 3.01E-04 | 2.18E-06 |
| 32 | GO:0005840 | ribosome | CELLULAR_COMPONENT | 5.00E-04 | 3.90E-06 |
| 33 | GO:0003824 | catalytic activity | MOLECULAR_FUNCTION | 9.90E-19 | 3.67E-21 |

Taken together, the good correlation of protein clusters with experimentally-validated sequences, the enrichment of expected annotation terms, and the presence of expected highly-abundant proteins or proteins critical to chloroplast biology suggest that both the RBH and UCLUST methods achieved good accuracy and sensitivity for genes with conserved chloroplast targeting which are likely critical in all photosynthetic plants for minimal chloroplast function. It is noteworthy that 194 clusters in RBH and 333 core clusters in UCLUST contain at least one Arabidopsis sequence but have no associated gene synonyms available (Additional Files 4 and 5). As the sensitivity for conserved plastid-targeted proteins was found to be very high overall, many of these 194-333 clusters with missing annotation information are likely biologically accurate, in which case they are excellent candidates for understanding hitherto uncharacterized aspects of chloroplast biology.

### Analysis of Semi-Conserved and Non-Conserved Plastid-Targeted Proteins

Semi-conserved plastid-targeted gene families in which predicted plastid-targeting was found for two or more sequences only in monocots, only in eudicots, or uniquely in *A. trichopoda* were identified beginning with the most diverse clades. In each case, all clusters with predicted plastid-targeted sequences or at least four predicted non-plastid-targeted sequences from the outgroup species were removed. A total of 572 gene families with plastid-targeted sequence specific to monocots and 430 to eudicots were found using RBH methods (Table 6, Figure 5), while UCLUST detected 1,054 and 885, respectively (Table 7, Figure 6). Additionally, 82 clusters with *Amborella*-specific plastid targeting were found using RBH, and 195 were found with UCLUST. These findings indicate that gene families with semi-conserved plastid-targeting outnumber core clusters by 73% in RBH and more than 150% in UCLUST. Narrowing focus to the subclade and family level revealed that semi-conserved clusters are still abundant, indicating that significant plastid proteome variation is present across all taxonomic levels. It is plausible that some of the clusters with plastid-targeting specific to either monocots or eudicots have functionally related clusters in the reciprocal group but lack sufficient homology to cluster together. Such an occurrence seems unlikely in most cases because the clustering methods used here were relatively liberal, but isolated cases may still occur. In some cases, non-orthologous or chimeric genes could also functionally replace an otherwise conserved gene and lead to loss of orthologous sequences in particular species or taxonomic groups [104, 105].

Finally, clusters with predicted plastid targeting only present in a single species were identified in RBH (Table 6, Figure 5) and UCLUST (Table 7, Figure 6). Singletons and clusters containing only a single genotype were discarded as these likely represent gene prediction errors. For example, predicted proteins in *Malus* which do not share homology with proteins in other species are typically poorly-supported by transcriptomics evidence: examination of over 300 such sequences revealed only one that had full coverage and was not a smaller fragment of a larger protein (data not shown). Since the chloroplast transit peptide is presumed to have arisen recently in each cluster, the term “nascent plastid-targeted proteins” (NPTPs) was coined to represent such proteins. Unsurprisingly, genotypes with large and complex genomes possessed a more significant number of NPTPs: *A. amnicola* had the least, at just 52 in RBH and 97 in UCLUST, while *P. virgatum* had the most, with 682 NPTPs found in RBH and 1,458 in UCLUST. The predicted proteome of *A. amnicola* is based on transcriptomics data rather than genome-wide prediction, while *P. virgatum* has the largest genome and most extensive predicted proteome of the species in this analysis, so these trends are consistent with expectations.

Additionally, up to 728 proteins were uniquely targeted to the plastid in *M.* × *domestica*, and between 300-400 proteins had species-specific plastid transit peptides in *B. distachyon*, *F. vesca*, *G. max*, *S. italica*, and *S. bicolor*. Arabidopsis had some of the lowest estimates of NPTPs, with only 74 found in RBH and 166 in UCLUST. Species-unique plastid-targeted proteins had a moderately linear correlation with the total number of sequences in each species R^2^=0.73 in RBH and 0.72 in UCLUST, Figure 7A), but the removal of the outlier *P. virgatum* resulted in nonlinear correlation (Figure 7B). Consequently, extreme increases in genome size and complexity are hypothesized to create more opportunities for the evolution of novel transit peptides and diversification of the plastid proteome, but differences are subtler when the genomes being compared are closer in size. Previous literature (e.g., [106–108]) has suggested that gene duplication is a prerequisite or at least greatly encourages neofunctionalization via novel subcellular targeting, and the generally linear correlation with proteome size suggests that this may indeed be the case. However, based on the data, the evolution of the plastid proteome is more likely to be driven by environmental adaptation and selection pressure [109].

**Figure 7:**
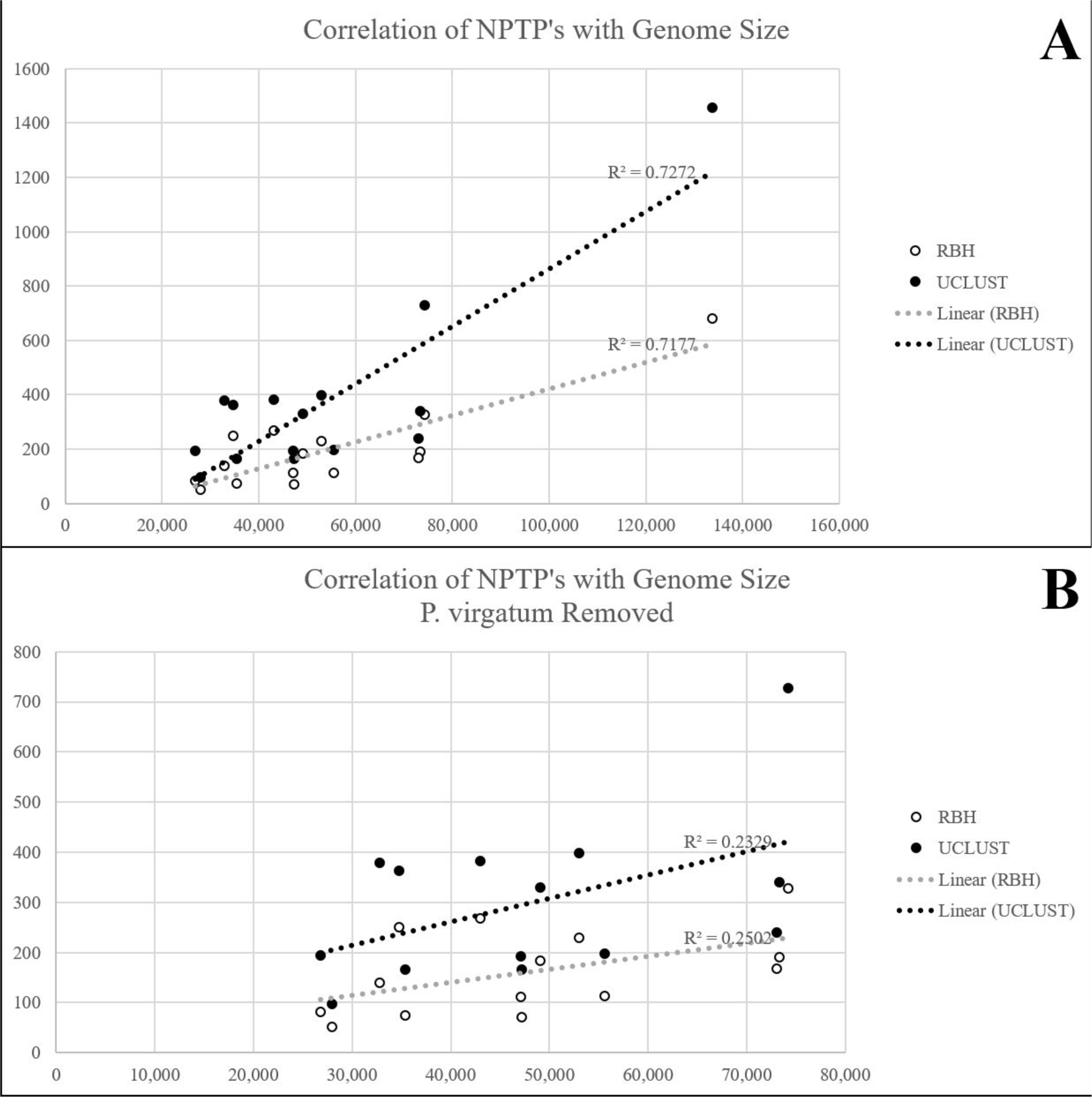
Correlation of Total Proteome Size with Nascent Plastid-Targeted Proteins (NPTPs). A. Clusters containing at least three species and with predicted plastid-targeted proteins in only one species were compared to the total proteome size for both RBH and UCLUST clustering methods. Although the correlation was moderately linear when *P. virgatum* was included, its extremely large proteome skewed results. **B**. Correlation after removal of *P. virgatum*. Weakly linear correlation indicates that the evolution of novel transit peptides is a random process.

As with the conserved plastid-targeted clusters, the accuracy of targeting prediction in NPTPs was cross-validated against experimentally-validated proteins from Arabidopsis. For the RBH clusters, 75% (4 proteins) were validated to be true plastid proteins via GFP, and 53.8% (17 proteins) validated by MS. For UCLUST, 29.4% (17 proteins) were validated by GFP, and 41.4% validated by MS. Specificity was also very high: only 6.3% of 300 predicted non-plastid-targeted proteins in RBH-generated NPTP clusters were found to actually be plastid-targeted by GFP, while the rate in MS-validated proteins was 13.4% (967 proteins). UCLUST generated similar results, with false negative error rates of 3% (493 proteins) in GFP-validated data and 12.5% (1,369 proteins) for MS-validated data. The few false negatives in predicted NPTPs may be representative of ambiguous/intermediate sequences in clusters which are already predicted to be uniquely chloroplast-targeted in Arabidopsis and therefore represent missing links. More pertinently, the GFP estimates are likely more accurate due to the experimental specificity errors inherent in mass spectrometry, and the 3-6% error rates are within an acceptable range.

Overall, these data affirm that evolution of the plastid proteome is highly dynamic at the species-level. Compared to previous reports, somewhat reduced species-unique plastid-targeted proteins are reported here (e.g., [2, 10]) due in part to the removal of singletons and single-species clusters. Homology to sequences in other species dramatically decreases the probability of pseudogenes and gene prediction errors. Remarkably, the monocot species had an average of 50-60% more species-unique plastid-targeted protein clusters than eudicot or *Amborella* counterparts. Even after removal of the outliers *P. virgatum* and *A. amnicola*, monocots still had 40% more plastid-targeted clusters than eudicots according to RBH methods, and over 80% more clusters using UCLUST. The reasons for this could be two-fold. First, the monocot species in this analysis have larger proteomes on average, increasing the overall likelihood for both *de novo* evolution of NPTPs and for retention of orphaned singleton/species-specific proteins. Secondly, monocots, and especially grasses, have been described to have many presence/absence variants (PAV’s) and copy number variants (CV’s) in their genomes. Pan-genome sequencing of *B. distachyon* revealed over 7,000 pan-genes that are not present in the reference genome, and an average of 9 Mb of sequence in each accession does not align to the reference genome [110]. Similar rates of PAV’s have been reported for cereal crops: only half of the pan-genome diversity of maize is present in the reference genome [111], over 21,000 predicted wheat genes are not represented in the reference genome [112], and 8,000 predicted rice genes are not represented in the Nipponbare reference genome [113]. In contrast, pan-genomes of Arabidopsis [114] and tomato [115, 116] describe variation primarily at the SNP and small insertion/deletion levels, although one report described that 14.9 Mb of the Columbia-0 genome was absent in one or more other accessions [117]. In *Brassica oleracea,* less than 20% of genes were affected by presence/absence variation [118]. Somewhat higher variation is observed in legumes: 302 soybean lines including varieties, landraces, and wild accessions revealed 1,614 copy number variants and 6,388 segmental deletions, and 51.4% of gene families were dispensable [119] while in *Medicago truncatula*, 67% of annotated genes may be dispensable [120]. It bears consideration that the pangenomes of the grasses are primarily within cultivated accessions and have already passed through a domestication filter which already significantly reduces genomic diversity, whereas the pangenomes of most of the eudicots include wild and landrace accessions.

These trends suggest that PAV’s and CV’s are significant drivers of plastid proteome evolution, either by retention of orphaned genes or by *de novo* evolution of transit peptides in duplicated genes. Despite the smaller number of species-unique clusters, conserved plastid-targeted proteins are still outnumbered up to 25-fold by species-unique or semi-conserved proteins. If even a fraction of these sequences is accurate and expressed *in vivo*, each could impart novel biological functions because escape from the evolutionarily established biochemical and regulatory environment could impart a different function in a new subcellular environment without changing the functional sequence of the protein. Thus, each of these is an excellent candidate for further characterization to determine if unique phenotypes are created by relocalization to the plastid.

Conversely, species-specific plastid-targeted genes in model systems could yield misleading interpretations because the same phenotypes for those genes would not be observed in species where homologs do not have plastid-specific localization. Such a situation is potentially problematic for the unique plastid-targeted proteins detected for Arabidopsis, *B. distachyon*, and rice because it is likely that some of these genes already have a described gene function that is being inaccurately ascribed to plants as a whole. Indeed, out of 113 Arabidopsis proteins with predicted species-specific plastid-targeting, 18 have a described phenotype, and 100 are cited in previous research reports (summarized in Additional File 3). In cases where the predicted localization divergence is validated, the mutant phenotypes for those sequences will have to be revised.

## Conclusions

The evaluations conducted in this study support the hypothesis that a combination of subcellular localization prediction programs can accurately predict chloroplast transit peptides at a whole-genome scale in higher plants and can perform equally well for both monocots and eudicots. The best-performing method was then applied to predict chloroplast proteins globally for a diverse range of angiosperm species and developed both a slow and accurate reciprocal best-BLAST hit method and a fast-liberal UCLUST method to cluster gene families. Though results were not identical, UCLUST yielded comparable results while performing more efficiently. With the addition of more genotypes, UCLUST could be a useful tool to overcome the inefficiency of BLAST-based methods. The consensus of both methods determined that the hypothesis of extreme plastid proteome variability was supported and robust across taxonomic space. Roughly 700 genes were shared between the chloroplast proteomes in all plant species, but these were vastly outnumbered by proteins with variable plastid targeting prediction. Most of these species- or clade-specific proteins have no known function for the plastid and are excellent candidates for further studies. Additionally, roughly a third of conserved plastid-targeted proteins have no known function and could be targeted for reverse genetics experiments in the future. Biological verification of these sequences remains a significant challenge. Even if good prediction accuracy was achieved, these sequences may be poorly expressed, expressed only in particular conditions, or are nonfunctional. Incorporation of transcriptomics would provide significant evidence that these genes are at least expressed, and patterns of gene expression along with co-expression information may also reveal additional information about their function. Experimental validation using mass spectrometry could also be used, but many proteins may have abundances below detection limits, and technical challenges also remain for the isolation of non-green plastids where they may be more abundant. The decreasing costs of gene synthesis make high-throughput fluorescence protein assays an attractive alternative. In addition to increased sensitivity and specificity compared with mass spectrometry, fluorescent protein assays could also be used to simultaneously validate whether the localization of species-unique proteins are truly different from their nearest predicted non-plastid-targeted homologs. The methods and results reported in this study will enable rapid, accurate and cost-effective identification of plastid-targeted proteomes in new plant species as their genomic information becomes available. These research findings are expected to provide a foundation for further research into unique plastid biology and to understand better how diversification of the organellar proteomes contributes to important agronomic, biochemical, culinary, or even aesthetic traits.

## Methods

### Cross-validation of in silico techniques

Test datasets for cross-comparison of subcellular prediction algorithms were retrieved from PPDB (2012 update; current as of this writing), AT_CHLORO (January 2015 update; current as of this writing) [23], Suba4 (30 June 2017 update; current as of this writing) [24], CropPAL version 58839ba [26], and CropPAL2 version 74866967 [26]. Headers which could not be referenced to the most up-to-date reference proteomes were discarded. For AT_CHLORO, Suba4, and PPDB databases, all genes located on the chloroplast and mitochondrial genomes were removed, and redundant headers were merged. Subsets of data including sequences confirmed by mass spectrometry, GFP fusion, either GFP or mass spectrometry, or both were extracted from each database by filtering for the keywords “Chloroplast” or “Plastid.” All ambiguous results containing experimental evidence for both plastids and at least one other subcellular fraction were removed.

Experimentally validated protein sequences were analyzed with TargetP v.1.1 [39, 40], WoLF PSORT Command Line Version 0.2 [36], PredSL Web Server [42], Localizer v.1.0.2 [37], MultiLoc2 version 2-26-10-2009 [48], and PCLR update 2011-11-24 release 0.9 [34]. Additionally, NLStradamus v.1.8 [121] was used as part of the Localizer algorithm, while Python v.2.7.5, LIBSVM v.2.8, BLAST v.2.2.30, and Interproscan v.5.25-64.0 were used as part of MultiLoc2. Results for each workflow were converted into binary classification and evaluated for Sensitivity (SE), Specificity (SP), Matthew’s Correlation Coefficient (MCC), and accuracy (ACC) as related to plastid localization prediction based on the number of true positives, false positives, true negatives, and false negatives compared to the annotations in the corresponding experimental dataset (see equations below). Combinatorial approaches were performed for each possible combination of programs from two up to all six programs, and different thresholds were evaluated based on the number of programs in agreement for plastid localization. Complete records of individual and combinatorial workflows for each experimental dataset are available in Additional File 2. The heatmap in Figure 1 was generated using conditional formatting in Microsoft Excel.

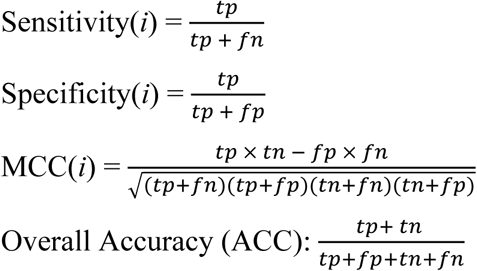

Where *tp* is the number of sequences correctly identified as plastid-targeted, *tn* is the number of sequences correctly predicted to be non-plastid-targeted, *fp* is the number of non-plastid-targeted sequences incorrectly predicted as plastid-targeted, and *fn* is the number of plastid-targeted sequences that were predicted as non-plastid-targeted. Note that these categorizations are based on the accuracy of the database annotation and any filtering that was applied to data subsets, and they may not reflect biological accuracy.

### Whole Proteome Analysis

Predicted proteomes for *Amborella trichopoda*, *Arabidopsis thaliana*, *Brachypodium distachyon*, *Fragaria vesca*, *Glycine max*, *Malus* × *domestica*, *Oryza sativa*, *Panicum virgatum*, *Populus trichocarpa*, *Prunus persica*, *Setaria italica*, *Solanum lycopersicum*, and *Sorghum bicolor* were downloaded from Phytozome [82]. The proteome of *Anthurium amnicola* was obtained by personal correspondence with Dr. Jon Suzuki in advance of the publication [71]. For *Vitis vinifera,* an expanded proteome version was obtained from [89]. For *Malus* × *domestica*, modifications to the predicted proteome were made because over 15,000 sequences, representing over 20% of the predicted proteome, were determined to have no significant matches to proteins from other species (See Additional File 3). The predicted proteome was expanded using apple transcriptomics data that were downloaded from the NCBI SRA database under the project numbers PRJEB2506, PRJEB4314, PRJEB6212, and PRJNA231737, representing a mixture of leaf, apical meristem, fruit, and root tissues at different time points and under varying conditions [77–80,122]. These sources are described further in Additional File 1. Sequence files were processed in CLC Genomics Workbench version 8 (Qiagen Bioinformatics, Hilden, Germany); paired Illumina read files and 454 sequencing files were indicated during import. Graphical QC reports were generated to obtain nucleotide contribution (GC content) and quality distribution (quality scores) by base position. Reads were processed to remove ambiguous nucleotides and base quality scores lower than 0.001. Illumina reads were additionally trimmed at the 5’ end until the GC content stabilized within 0.5% of the average, and reads with fewer than 34 bases remaining were discarded. All paired read files were subsequently merged using default settings. All processed read files were assembled *de novo* with default settings. Assembled contigs of >300 bp were kept and used to predict open reading frames (ORF’s). Non-overlapping ORF’s with at least 5x average base coverage and >300 bp were extracted and translated into protein sequences. Finally, extracted protein sequences were compared against the existing *Malus* × *domestica* v.1.0 predicted gene set [76] downloaded from Rosaceae.org. All hits with greater than 98% ID and coverage (as per [80]) were tagged as potential duplications or alleles of the original headers but were kept in the peptide dataset in case minor mutations caused differential localization prediction. All sequences generated in this transcriptomics experiment are available in Additional File 6. In total, 36,477 sequences were obtained, of which 26,881 sequences were determined to be unique in comparison with the apple genome paper [76]. Addition of the unique genes from the *de novo* transcriptome created a final dataset of 64,680 unique proteins. Redundant sequences from the resulting transcriptome were retained in case minor differences resulted in differential targeting.

The predicted proteomes of all species were filtered to remove any sequences less than 100 residues and which did not begin with methionine. Post-analysis filtering was accomplished by removing singleton sequences that failed to find matches with both the USEARCH method and BLAST (indicated for each sequence in Additional File 3). Remaining sequences were analyzed with TargetP v.1.1 [39, 40] and Localizer v.1.0.2 [37]. All sequences predicted by both methods to have a chloroplast transit peptide were classified as plastid-targeted, and all sequences with either “1 or 2” or “0 of 2” chloroplast transit peptide predictions were classified as non-plastid-targeted.

### Clustering of Gene Families

Reciprocal Best-BLAST hit clustering was performed as follows: Pairwise BLAST-P (v.2.3.0+ command line executable; [123, 124]) was performed for each species’ predicted proteome set against that of every other species in both forward and reverse directions. These results were filtered for hits in which identity and coverage parameters exceeded 40%. Of these, only hits in which two sequences from different genomes were the respective best hit were kept. Next, better-BLAST hits within each species were performed by conducting pairwise BLAST-P of the predicted proteome against itself. Hits exceeding 90% coverage and identity and which was reciprocal within the first 10 hits were collected. Cluster merging was performed by iterating through each possible header and collapsing all pairwise hits containing that header.

Clustering using the UCLUST algorithm proceeded as follows: An initial run on a length-sorted FASTA file containing all sequences was performed using ‘Cluster_Fast’ function of UCLUST (v.9.2.64_win32; [125]) with 40% identity and 40% query coverage. Next, random seeds were constructed by extracting a single random sequence from each cluster, sorting the resulting sequences by length, and appending them to a length-sorted FASTA of the full sequence list used in the initial “Cluster_Fast” analysis. 100 randomly-seeded FASTA files were then analyzed with “Cluster_Fast” set to 90% sequence identity. Target and query coverage were additionally set to 0.4 to avoid problems with small query sequences acting as centroids for much larger sequences as a result of USEARCH being performed in sequential rather than length-sorted order. Cluster merging was performed by iteratively searching through each possible sequence header and collapsing all clusters containing that header. Custom scripts were developed for automating program workflows, referencing and translating sequences or headers, performing seed randomization for the modified UCLUST technique, performing cluster expansion, calculating statistics on clustering outputs, and referencing headers to respective clusters for both workflows. Sequence members within merged clusters from RBH and UCLUST methods were referenced to the predicted plastid targeting phenotype, and all clusters containing plastid-targeted members were extracted. Conserved plastid-targeted gene families were defined as clusters containing at least 13 species and in which all had either predicted plastid transit peptides or at least three additional sequences. Semi-conserved plastid-targeted gene families were defined as clusters containing plastid-targeted sequences from at least 2 species within each family or clade and no predicted plastid-targeted sequences from species outside that clade. Non-conserved plastid-targeted gene families were defined as all clusters containing a minimum of three species in which only one species had a plastid-targeted sequence.

### Gene Ontology Enrichment

Annotations for NPTPs were retrieved from Phytozome [82] for each of the species used in the analysis except *Anthurium amnicola* and *Vitis vinifera*, which were retrieved from [71] and [89], respectively. Non-redundant predicted proteins produced by the *de novo* transcriptome assembly of *Malus* × *domestica* were annotated using BLASTP against the NR Protein database at NCBI with BLAST2GO v.4.1.9 default parameters [103] (BioBam Bioinformatics, Valencia, Spain). GO terms were converted into GOslim annotations using BLAST2GO, and for each cluster, all terms shared by at least three species and present in over 10% of a cluster’s sequences were extracted to develop query datasets. In parallel, the same methods were used to extract GO terms from the total list of clusters to serve as reference datasets. Enrichment of GO terms in the shared plastid-targeted clusters was performed using BLAST2GO, with Fisher’s Exact Test was used to calculate significance using a false discovery rate (FDR) of less than 0.05 as a minimum significance threshold [103]. Graphical analyses of enriched GO terms were produced in BLAST2GO.

### Gene and Phenotype Identification

Full gene annotations include described gene names were downloaded for the TAIR10 Arabidopsis genome from Phytozome [82]. Gene names were referenced from the annotation file for Arabidopsis sequences present in conserved plastid-targeted protein clusters. Phenotype information for species-unique plastid-targeted proteins was referenced on NCBI [126].

## Supporting information

Supplemental Plastid Clustering

## Authors’ contributions

RWC and AD designed the study. RWC performed localization prediction, gene clustering, and data analysis. EHR assisted in methods development. SLH performed gene annotation analyses. AD and EHR supervised the study. RWC and AD prepared the manuscript. All authors read and approved the manuscript. The authors declare no conflict of interest.

## Availability of data and materials

“The datasets supporting the conclusions of this article are included within the article and its additional files. Perl scripts used in the organization of data and execution of protein clustering are available at Sourceforge under the Project Name “Plastid Variation” and the homepage https://sourceforge.net/p/plastid-variation.

Operating System(s): Platform Independent

Programming Language: Perl Other

Requirements: TargetP v.1.1, Localizer v.1.0.2, BLAST v.2.3.9+ command line executable, UCLUST v.9.2.64_win32, RAxML v.8.2.31, MUSCLE v.3.8.31, MAFFT v. 7.407, Phyutility v2.2.6, FastTree 2.1.10.

License: open source

Restrictions for use by non-academics: no restrictions.

## Acknowledgments

Work in the Dhingra lab was supported by Washington State University Agriculture Center Research Hatch Grant WNP00011 to AD. RC and SH acknowledge the support received from the National Institutes of Health/National Institute of General Medical Sciences through an institutional training grant award T32-GM008336. The contents of this work are solely the responsibility of the authors and do not necessarily represent the official views of the NIGMS or NIH.

